# Calcium-dependent protein kinase 5 links calcium-signaling with *SARD1*-dependent immune memory in systemic acquired resistance

**DOI:** 10.1101/553750

**Authors:** Tiziana Guerra, Silke Schilling, Fabian-Philipp Sylvester, Benjamin Conrads, Tina Romeis

## Abstract

- Systemic acquired resistance (SAR) prepares infected plants for faster and stronger defense activation upon subsequent attacks. SAR requires an information relay from primary infection to distal tissue and the initiation and maintenance of a self-maintaining phytohormone salicylic acid (SA)-defense loop.
- In spatial and temporal resolution we show that calcium-dependent protein kinase CPK5 contributes to immunity and SAR. In local basal resistance CPK5 functions upstream of SA-synthesis, -perception, and -signaling. In systemic tissue, enhanced CPK5 signaling leads to an accumulation of SAR marker genes including transcription factor *Systemic Acquired Resistance Deficient 1* (*SARD1*).
- Plants of enhanced CPK5-, but not CPK6-, signaling display a ‘super-priming’ phenotype of enhanced resistance toward a secondary bacterial infection. In *sard1* background, CPK5-mediated basal resistance is still mounted but systemic ‘super-priming’ is lost.
- The biochemical analysis determines CPK5 half maximal kinase activity for calcium K50 [Ca^2+^] to ∼100 nM close to the cytoplasmic resting level. This low activation threshold uniquely qualifies CPK5 to decode subtle changes in calcium prerequisite to immune signal relay and to onset and maintenance of priming at later time points in distal tissue. Our data explain why CPK5 functions as a hub in basal and systemic plant immunity.

## Introduction

Rapid and long-term activation of the plant immune system guarantees plant survival upon pathogen infection. Pathogen recognition initiates early intracellular responses at the local infection site, which involve changes in ion fluxes inclusive an increase of the cytoplasmic calcium concentration, recruitment of signaling cascades and the activation of transcriptional reprogramming. The information of local ‘having been attacked’ is subsequently relayed to distal plant parts. On the basis of phytohormone-mediated transcriptional and metabolic changes, resistance to the attacking pathogen is manifested in the entire plant. The plant may establish and maintain systemic acquired resistance (SAR) and thus prime an immune memory of ‘being prepared to defend upon a subsequent attack’ (Hake & Romeis, 2018).

During defense initiation, a microbial pathogen is recognized at the site of infection via non-species specific Pathogen-Associated Molecular Patterns (PAMPs). PAMPs bind as ligands to specific receptors (Pattern Recognition Receptors (PRRs)), intracellular defense responses become activated and PAMP-triggered immunity (PTI) is established. In a species-specific context, a second layer of defense, Effector-Triggered Immunity (ETI), has evolved to detect PTI-suppressing effectors from adapted pathogens. In ETI defense response activation occurs either directly upon effector binding, or indirectly by the recognition of a process of effector target modification. ETI triggers a stronger more long-term response and hypersensitive cell death reaction. Also, ETI may result in ‘priming’ of SAR rendering a plant prepared for repeated infection after a time gap. Primed plants show more rapid and stronger activation of defense responses upon a secondary infection even by unrelated pathogens (Conrath, 2006; Hilker *et al.*, 2015; Hake & Romeis, 2018).

Both, local and systemic resistance depend on the synthesis, accumulation, and downstream signaling of the phenolic phytohormone salicylic acid (SA). SA biosynthesis is catalyzed by ISOCHORISMATE SYNTHASE 1 (ICS1) and SA transport from the chloroplast to the cytoplasm is mediated by ENHANCED DISEASE SUSCEPTIBILITY 5 (EDS5) (Wildermuth *et* al., 2001; Serrano *et al.*, 2013). SA-dependent responses were shown to be under the control of PHYTOALEXIN DEFICIENT 4 (PAD4). PAD4, in concerted action with a lipase-like protein, ENHANCED DISEASE SUSCEPTIBILITY 1 (EDS1), is required for the expression of multiple defense responses in basal defense in PTI and ETI as well as for ETI-associated hypersensitive response leading to systemic resistance (Glazebrook *et al.*, 1997; Wiermer *et* al., 2005; Rietz *et al.*, 2011). SA is perceived by NON-EXPRESSOR OF PATHOGENESIS-RELATED GENES 1 (NPR1) as one of three known SA-receptors (Fu *et al.*, 2012; Wu *et al.*, 2012; Yan & Dong, 2014; Manohar *et al.*, 2015; Y. Ding *et al.*, 2018). Continuous SA-signaling will thus impact on changes in histone modification, in gene expression, and in metabolite production and lead to persistent plant resistance to pathogens.

This genetic framework subsequent to local pathogen attack in SA-mediated plant resistance is well characterized. Yet little is known about the switch into prolonged systemic continuous SA signaling and the establishment of SAR, a decision which comes for a plant along with a cost on development and severe growth retardation (Hake & Romeis, 2018).

The nature of plant signals that bear the information of ‘having been attacked’ from local infection sites to distal tissues is an ongoing matter of debate. Several metabolites have been described, whose requirement for distal SA accumulation and SAR have been proven, including non-proteinaceous amino acid pipecolic acid (Pip) (Návarová *et al.*, 2012) and most recently N-hydroxypipecolic acid (NHP) (Hartmann *et al.*, 2018). Pip has been recognized as a most prominent signal molecule and marker for SAR. It is synthesized 12 h post pathogen infection from L-Lys through enzymes AGD2-LIKE DEFENSE RESPONSE PROTEIN 1 (ALD1) and SAR-DEFICIENT 4 (SARD4) (Bernsdorff *et al.*, 2016; P. Ding *et al.*, 2016; Hartmann *et al.*, 2017). Pip is further modified through FLAVIN-DEPENDENT MONOOXYGENASE 1 (FMO1) to generate NHP. Remarkably, Pip enforces a prolonged SA biosynthesis in systemic plant tissue in an *ALD1*/*FMO1*/*ICS1* positive feed-forward loop during SAR.

The manifestation of SAR requires key transcriptional regulator SYSTEMIC ACQUIRED RESISTANCE DEFICIENT 1 (SARD1) which is induced late after a priming pathogen attack (Wang *et al.*, 2009; Truman & Glazebrook, 2012; Sun *et al.*, 2015). In distal plant tissue SARD1 mediates continuous SA and Pip synthesis. *sard1* mutants no longer establish SAR (Zhang *et al.*, 2010; Wang *et al.*, 2011) and thus lack the memory of ‘having been attacked’. Calcium-Dependent Protein Kinases (CDPKs, in *Arabidopsis thaliana*: CPKs) are calcium-sensor - protein kinase effector proteins in a single molecule. Upon calcium binding, a consensus enzyme with four EF-hand calcium-binding motifs undergoes a conformational change, adopts a kinase-active state and thus phosphorylates target proteins (Cheng *et al.*, 2002; Liese & Romeis, 2013; Schulz *et al.*, 2013). CDPKs have consequently been discussed as decoders of changes in the cytoplasmic calcium concentration [Ca^2+^]. Accordingly, CDPKs were identified as positive regulators during the initiation of immune signaling, triggering local immune responses in PTI and ETI, and contributing to basal resistance (Kobayashi *et al.*, 2007; Boudsocq *et al.*, 2010; Kobayashi *et al.*, 2012; Dubiella *et* al., 2013; Gao *et al.*, 2013; Gao & He, 2013). A negative regulatory role in immune signal initiation was demonstrated for CPK28 (Monaghan *et al.*, 2014; Monaghan *et al.*, 2015; J. Wang *et al.*, 2018).

Furthermore, CDPKs are implicated in the relay of an immune signal from local to distal sites. In particular, Arabidopsis CPK5 has been implicated to drive a calcium- and ROS-based auto-propagating signaling loop, which contributes to signal propagation from a local infection site to un-infected foliar tissue of a plant in PTI (Dubiella *et al.*, 2013; Seybold *et al.*, 2014; Hake & Romeis, 2018).

However, a specific requirement of calcium signal decoding during the manifestation of SAR in systemic plant tissue at later time points after a primary local pathogen attack is unknown. Plants overexpressing CPK5-YFP show constitutive CPK5 enzyme activity. Enhanced CPK5–signaling leads to constitutive defense responses and increased SA-dependent disease resistance in these plants (Dubiella *et al.*, 2013). Independently, several alleles of *cpk5* have been identified in a forward genetic screen as suppressors of the autoimmune mutant *exo70B1* characterized by its resistance to multiple bacterial and fungal pathogens. *exo70B1* plants display constitutive SA-dependent defense responses reminiscent to those of enhanced CPK5-signaling. Resistance depends on the atypical immune receptor TN2, and TN2 protein interacts with CPK5 stabilizing the enzyme in a kinase-active state (Liu *et al.*, 2017). Taken together, these data suggest that CPK5 contributes to the control of long-term systemic resistance and SAR.

Here, we address in temporal and spatial resolution the role of CPK5 as a calcium-regulated key component for signaling in systemic resistance. CPK5 functions upstream of the SA signaling cascade comprised by *ICS1, EDS5*, and *NPR1* as well as by *PAD4*, and mediates *ALD1*- and *FMO1*-dependent immune responses in basal and systemic resistance. Furthermore, enhanced CPK5 signaling, but not CPK6 signaling, causes an increase in SAR at late time points in distal tissue, and this ‘super-priming’ requires *SARD1*. *sard1* mutant plants still display CPK5-mediated enhanced basal resistance but lack the memory phase to gain ‘super-priming’. CPK5 is capable of responding to even subtle calcium changes due to its low biochemical K50 for calcium at ∼100 nM. This feature qualifies predominantly CPK5 over CPK6 an ideal component in signal propagation and in the systemic control of defense manifestation through a switch in *SARD1*-dependent SAR.

## Material and Methods

### Plant materials and growth conditions

*Arabidopsis thaliana* Col-0 wild type, mutants, CPK5- and CPK6-overexpressing lines and derived crosses were grown under short day conditions (8 h light per 16 h dark cycle) at 20°C, 60% relative humidity. The mutant plants represent *cpk5* (SAIL_657C06) and *cpk6* (SALK_025460C). *CPK5-YFP#7* was crossed to *ics1* (SALK_088254), *npr1-1* (N3726), *pad4* (SALK_089936), *eds5* (SALK_091541C), *sard1-1* (SALK_138476C), *ald1* (SALK_007673), and *fmo1* (SALK_026163) and subsequently genotyped in filial generations. Primers used for genotyping are listed in Supplementary Table 1. Plant lines *ics1, pad4, eds5*, and *sard1* were obtained from the Nottingham Arabidopsis Stock Centre (NASC), *npr1-1* was kindly provided by C. Gatz (University Göttingen, Germany), *ald1* and *fmo1* were kindly provided by J. Zeier (University Düsseldorf, Germany). *CPK5-YFP#7* and CPK5mut-YFP #15 (expressing a kinase-deficient variant of CPK5) have been described previously (Dubiella *et al.*, 2013).

### Generation of CPK6-YFP overexpressing line

The coding region of full-length CPK6 (AT2G17290) was amplified from Col-0 cDNA with primers CPK6-YFP-LP and CPK6-YFP-RP. The 1636 bp CPK6 fragment was cloned into the pENTR D/TOPO vector (Invitrogen) and confirmed by sequencing. The C-terminal YFP-fusion construct was generated by LR-Gateway recombination into pXCSG-YFP. Transgenic *Arabidopsis* plants were generated by the floral dip method. The flowering *Arabidopsis thaliana* Col-0 plants were dipped into *Agrobacterium tumefaciens* GV3101 pMP90RK carrying pXCSG-CPK6-YFP. Seeds were harvested and selected for BASTA-resistance to gain independent transformants.

### Generation of CPK5-StrepII, CPK6-StrepII and CPK5-YFP-StrepII constructs

The C-terminal CPK6 StrepII-fusion construct was generated by using the pENTR D/TOPO CPK6 vector via LR-Gateway recombination into pXCSG-StrepII. The coding region of full-length CPK5 (AT4G35310) and CPK6 were amplified from pXCSG-CPK5-StrepII, pXCS-CPK5-YFP (Dubiella *et al.*, 2013) and pXCSG-CPK6-StrepII with primers CPK5-StrepII-LP and CPK5-StrepII-RP, primers CPK5-YFP-StrepII-LP and CPK5-YFP-StrepII-RP, and CPK6-StrepII-LP and CPK6-StrepII-RP, respectively. The 1749 bp CPK5-StrepII and 1707 bp CPK6-StrepII fragments were cloned into pET30 expression vector derivate (Novagen, Merck) denominated pET30-CTH (Glinski *et al.*, 2003) via *NdeI* and *XhoI* restriction sites to replace His-tag. To generate pET30-CPK5-YFP-StrepII a fragment encompassing 264 bp of CPK5 (C-terminal part) plus 720 bp YFP was cloned into pET30-CPK5-StrepII using the internal restrictions sites *MfeI* and *BsaI*.

### Gene expression analysis by RT-qPCR

To analyze transcript levels RNA was extracted from leaf tissue using the TRIzol method (ThermoFisher). 2 μg of RNA was incubated with RNase-free DNase (Fermentas) and subjected to reverse transcription using SuperscriptIII SuperMix (ThermoFisher) according to the manufacturer’s protocols. Real-time quantitative PCR analysis was performed in a final volume of 10 μl according to the instructions of Power SYBR Green PCR Master Mix (Applied Biosystems) using the CFX96 system (Bio-Rad). Amplification specificity was evaluated by post-amplification dissociation curves. *ACTIN2* (At3g18780) was used as the internal control for quantification of gene expression. Primer sequences are listed in Table S2.

### Protein expression and purification

The expression vector pET30 containing CPK5-YFP-StrepII, CPK5-StrepII or CPK6-StrepII was introduced in *E. coli* BL21 (DE3) (Stratagene) strain. Bacteria were grown at 37 °C in LB media containing 50 µg/ml kanamycin and protein expression was induced at OD_600_ 0.5-0.8 with 0.3 mM isopropylthiol-β-galactoside (IPTG) and incubated for an additional 2-2.5 h at 28 °C. Cells were harvested by centrifugation. Cells were lysed in lysis buffer (100 mM TRIS pH 8.0, 150 mM NaCl, 20 mM DTT, 100 µg/ml Avidin and 50 µl protease inhibitor cocktail for histidine tagged proteins per 1 g *E. coli*) by incubation for 20 minutes with 1 mg/ml Lysozyme and additional sonification. Cell debris was removed by centrifugation. The supernatant was loaded on self-packed columns containing 1 ml Strep-Tactin-Macroprep (50% suspension, IBA) equilibrated with wash buffer (100 mM TRIS pH 8.0, 150 mM NaCl). Flow-through was loaded on columns three times. After washing columns with 3 ml wash buffer, proteins were eluted with 3x 500 µl elution buffer (100 mM TRIS pH 8.0, 150 mM NaCl, 2 mM d-Desthiobiotin). Protein purity and concentration were analyzed via Bradford (Protein assay, Bio-Rad) and 10 % SDS-PAGE and Coomassie staining.

### *In vitro* kinase assay

*In vitro* kinase assay was performed with StrepII-tag affinity purified recombinant CPK5-variants and CPK6 with substrate peptides Syntide 2 or RBOHD (aa 141-150, encompassing S148: RELRRVFSRR). The final kinase reaction (30 µl) contained ∼25 nM CPK5-variants or CPK6 in a volume of 5 µl and either 20 µl buffer E (50 mM TRIS-HCl pH 8.0, 2 mM DTT, 0.1 mM EDTA) and 5 µl reaction buffer (60 mM MgCl_2_, 60 µM RBOHD S148 or 60 µM Syntide 2, 60 µM ATP, 3 µCi [γ-^32^P] ATP, 0.2 x buffer E, 30 µM CaCl_2_ or 12 mM EGTA) for time course analysis or 20 µl calcium buffer (30 mM TRIS pH 8.0, 10 mM EGTA, 150 mM NaCl and different concentrations of CaCl_2_ as indicated) and 5 µl reaction buffer without additional CaCl_2_ or EGTA for K50 determination. The reaction was either incubated for 20 minutes at 22°C for K50 determination or as indicated for time course analysis. The reaction was stopped by adding 3 µl 10 % phosphoric acid. Radioactive labeled phosphorylation of peptides was determined by the P81 filter-binding method as described (Romeis *et al.* (2000)). Kinase activities are plotted against the Ca^2+^ concentration using graph GraphPad Prism 4 software (GraphPad Software) in a four parameter logistic equation. Relative kinase activities in percentage are fitted by a four parameter logistic equation with the top fixed to 100% constant value using graph GraphPad Prism 4.

### In-gel kinase assay

Phosphorylation events were monitored in unstressed *Arabidopsis* two-week-old seedlings as described previously (Dubiella *et al.*, 2013). Seedlings were grown on 0.5 MS + 1% sucrose for 2 weeks and harvested in pools of 200 mg and directly frozen in liquid nitrogen.

### Bacterial growth *in planta*

To quantify bacterial growth in *Arabidopsis*, six-week-old plants grown on soil under short day conditions were used. The different *Pseudomonas syringae* strains were grown in King’s B media overnight at 28°C in a shaker at 200 rpm. The cells were harvested by centrifugation at 4°C for 15 min at 2,000xg and washed twice with 10 mM MgCl_2_. Cells were resolved in 10 mM MgCl_2_ to a concentration of 10^4^ cfu/ml used for infiltration into the leaf. For basal resistance *Pseudomonas syringae pv. tomato* DC3000 was used. In the case of SAR experiments, the primary infection and treatment with either bacterial suspension at OD_600_ 0.005 (avirulent *Pseudomonas syringae pv. maculicola* ES4326 *avr Rpm1*) or the control of 10 mM MgCl_2_ were conducted for two days in three fully developed ‘primary’ leaves, which were cut 24 h after infiltrations. For the secondary infection the virulent strain *Pseudomonas syringae pv. maculicola* ES4326 was used. Samples were harvested at day 0 and day 3 post (the secondary) infection. Per *Arabidopsis* line at least 8 plants were inoculated. For the analysis of systemic signal propagation, six-week-old *Arabidopsis* plants were used. Experiments were performed as described previously (Dubiella *et al.*, 2013).

### Gas chromatographic analysis of amino acid derivatives

The analysis was performed as described by Návarová *et al.* (2012). Statistical analysis was done using student’s t-test.

### Accession numbers

Sequence data from this article can be found in the Arabidopsis Information Resource or GenBank/EMBL databases under the following accession numbers: *CPK5* (AT4G35310), CPK6 (AT2G17290), *ICS1* (AT1G74710), *NPR1* (AT1G64280), *PAD4* (AT3G52430), *EDS5* (AT4G39030), *SARD1* (AT1G73805), *ALD1* (AT2G13810), *FMO1* (AT1G19250).

## Results

### SA biosynthesis and signaling are required for CPK5-mediated defense marker gene expression and enhanced pathogen resistance

To address whether SA synthesis and SA signaling are required for CPK5-mediated enhanced resistance to bacterial pathogens we performed crosses between the CPK5-overexpressing line *CPK5-YFP#7* and defense mutants *ics1, eds5, npr1-1, pad4*, and *sard1*. The phenotypic analysis revealed that reduced rosette diameter, necrosis, and crinkled leaves, which are attributed to enhanced CPK5-activity in line #7, are no longer evident in crossing lines in the absence of SA biosynthesis (*ics1*), transport (*eds5*) or signaling (*npr1*) (Fig. **1a**). This phenotypic reversion is corroborated by the expression of *PR1*, indicative for SA signaling, and *FRK1*, an flg22-responsive marker gene. *CPK5-YFP#7* is characterized by constitutive high levels of *PR1* and *FRK1* (Dubiella *et al.*, 2013), whose basal gene expression is reverted to Col-0 levels in the crossing lines (Fig. **1b**). Accordingly, in pathogen growth assays with the virulent pathogen *Pseudomonas syringae pv. tom*ato (*Pst*) DC3000 enhanced bacterial resistance indicative for *CPK5-YFP#7* is compromised, and crossing lines show either bacterial counts as the Col-0 wild type (x *npr1*) or become even more susceptible than Col-0 (x *ics1,* x *eds5*) (Fig. **1c**). The expression and presence of active CPK5-YFP protein kinase was confirmed by western blot and in-gel kinase assay as exemplarily shown for crossing lines to *ics1* and *npr1* (Fig. **S1a,b**). These data indicate that SA biosynthesis and SA downstream signaling are predominantly responsible for CPK5-mediated defense reactions.

**Fig. 1:**
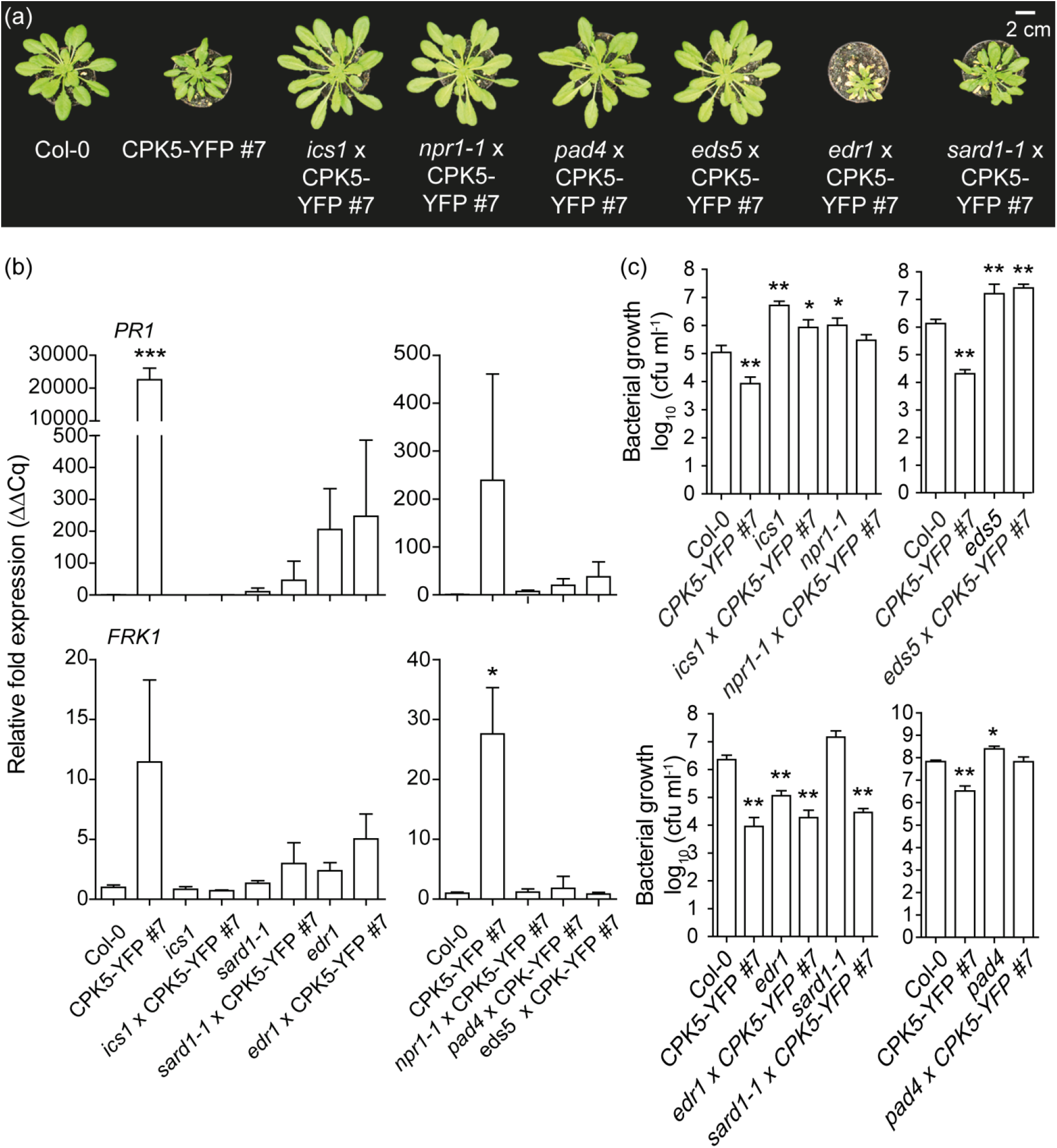
CPK5 signaling-dependent basal pathogen resistance requires SA-biosynthesis and signaling but is independent on *SARD1*. (**a**) Six-week-old plants of Col-0, *CPK5-YFP#7*, and of derived crosses with *ics1, npr1, pad4, eds5, edr1, and sard1*. Scale = 2 cm. (**b**) Basal *PR1* and *FRK1* gene expression of plants as shown in (a) was analyzed by RT-qPCR. Bars represent mean value ± SEM of three biological replicates. The asterisks indicate statistically significant differences in comparison to Col-0 (one-way ANOVA; Dunnett posttest; ***, *P*<0.001; *, *P*<0.05). (**c**) Six-week-old plants as shown in (a) were inoculated with *Pseudomonas syringae pv. tomato* (*Pst*) DC3000. Bacterial numbers were monitored 0 dpi (data not shown) and 3 dpi. Bars represent mean value ± SEM of 12 biological replicates. The asterisks indicate statistically significant differences in comparison to Col-0 (one-way ANOVA; Dunnett posttest; **, *P*<0.01; *, *P*<0.05). The experiment was repeated three times with similar results.

Likewise, *PAD4*, a key component and positive regulator of defense responses in the context of ETI upon activation of TIR-NB-LRR proteins, is required for CPK5-dependent and SA-mediated bacterial resistance. In line *pad4* x CPK5-YFP#7, the plant rosette diameter, defense marker gene expression, and pathogen growth are reverted to Col-0 phenotypes (Fig. **1**). The crossing line *edr1* x *CPK5-YFP#7* shows reduced *PR1* and *FRK1* defense expression compared to *CPK5-YFP#7*, but still elevated compared to Col-0, and retains its increased higher basal resistance, accompanied by the phenotype of a small plant rosette and the development of necrosis symptoms. Compared to an increased susceptibility of *ics1* and *npr1* single mutants crossing lines show a certain benefit of enhanced CPK5-signaling resulting in less bacterial growth (Fig. **1b**). These data hint toward an additional contribution of CPK5-triggered defense activation independent of *ICS1* and *NPR1*.

### The expression of SAR marker genes depends on CPK5

We next assessed SAR marker gene expression *ALD1* and *FMO1* in a temporally and spatially defined manner using a modified systemic signaling assay. Three local (proximal) leaves were stimulated by either mock-(10 mM MgCl_2_) or 200 nM flg22-infiltration. Samples of systemic tissue were harvested two days after the local infiltration (2 dpi) and gene expression was monitored via RT-qPCR. Enhanced CPK5 signaling in *CPK5-YFP#7* resulted in high constitutive *ALD1* and *FMO1* expression, irrespectively of a mock or flg22 treatment (Fig. **2a**). Likewise, *NHL10*, a marker gene for rapid CPK5-dependent defense signal propagation, is constitutively expressed in systemic tissue. In contrast, Col-0 and *CPK6-YFP#23* do not display elevated SAR marker gene expression. Single mutant lines of *cpk5* more than of *cpk6* show some reduction in basal and systemic *ALD1, FMO1* and *NHL10* expression at 2 dpi even after the local flg22 stimulus (Fig. **2b**). These data corroborate a specific role of CPK5 over CPK6 in the regulation of temporal late and spatially distal expression of defense genes.

**Fig. 2:**
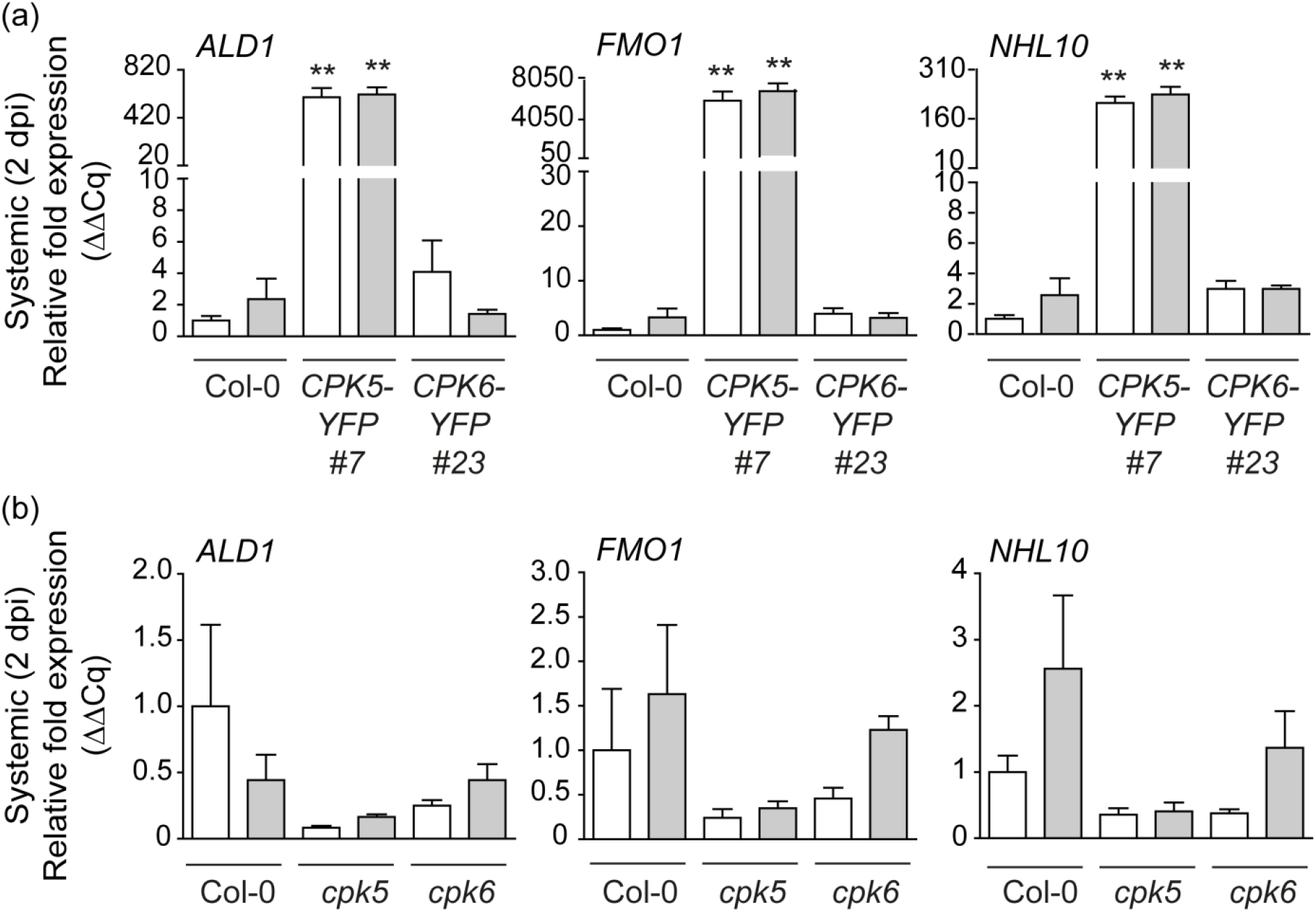
CPK5 but not CPK6 is involved in late systemic defense signaling. (**a**) Enhanced CPK5-signaling results in constitutive systemic expression of *ALD1, FMO1,* and *NHL10* at 2 dpi. Three local leaves of six-week-old plants of Col-0, *CPK5-YFP#7*, and *CPK6-YFP#23* were infiltrated with 10 mM MgCl_2_ (mock/white bars) or 200 nM flg22 (grey bars). After two days, three systemic leaves were harvested and gene expression was analyzed by qRT-PCR. Bars represent mean value ± SEM of three biological replicates. The asterisks indicate statistically significant differences in comparison to Col-0 mock (one-way ANOVA; Dunnett posttest; **, *P*<0.01). The experiment was repeated with similar results. (**b**) Systemic defense gene expression at 2 dpi is reduced in *cpk5*. Col-0, *cpk5*, and *cpk6* leaves were analyzed as described in (a). Bars represent mean value ± SEM of three biological replicates. The experiment was repeated with similar results.

### CPK5 contributes to priming

To address whether the observed constitutive *ALD1* gene expression correlates with SAR marker metabolite Pip accumulation we determined Pip levels by gas chromatography-mass spectrometry (GC-MS) analysis in Col-0 and *CPK5-YFP#7* plants. A statistically significant accumulation of Pip was measured in CPK5-overexpressing lines in the absence of pathogen exposure (Fig. **S2**).

Next, we conducted SAR bacterial growth assays where plants were exposed to a priming infection with avirulent *Pseudomonas syringae pv. maculicola* ES 4326 avrRpm1 (*Psm avrRpm1*) in local (proximal) leaves, followed two days later by a triggering infection with virulent *Psm* ES 4326 in distal leaves (Fig. **3** and scheme in Fig. **S3**). The reported benefit of priming as reduced bacterial growth upon a secondary infection is observed in Col-0 plants (Fig. **3**, Fig. **4d**, Fig. **5b**) (Durrant & Dong, 2004; Mishina & Zeier, 2006; Fu & Dong, 2013; Gruner *et al.*, 2013; Shah & Zeier, 2013). CPK5 overexpression leads to an overall increased resistance towards bacterial pathogens due to CPK5-mediated, SA-dependent enhanced basal resistance (Dubiella *et al.*, 2013) (Fig. **1c**, Fig. **3b**, Fig. **4c**,**d**, Fig. **5b**). Interestingly, this basal level of CPK5-mediated resistance, already seen upon infection with the virulent strain, mimics the SAR level in primed Col-0 plants. Remarkably, when *CPK5-YFP#7* is subjected to a combination of priming and triggering infections, a status of hyper-resistance (‘super-priming’) is observed (Fig. **3b**, Fig. **4d**, Fig. **5b**). No alteration in priming capacity compared to Col-0 is observed in plants overexpressing CPK6 (*CPK6-YFP#23*) (Fig. **3d**). A loss of priming capability is observed in *cpk5 cpk6* double mutant lines (Fig. **3e**), whereas no statistically significant reduction of priming occurs in respective *cpk5* and *cpk6* single mutant lines (Fig. **3a,c,e**). These data are consistent with previous reports in which single *cpk* mutants did not show compromised resistance to infections with virulent bacterial pathogens (Boudsocq *et al.*, 2010).

**Fig. 3:**
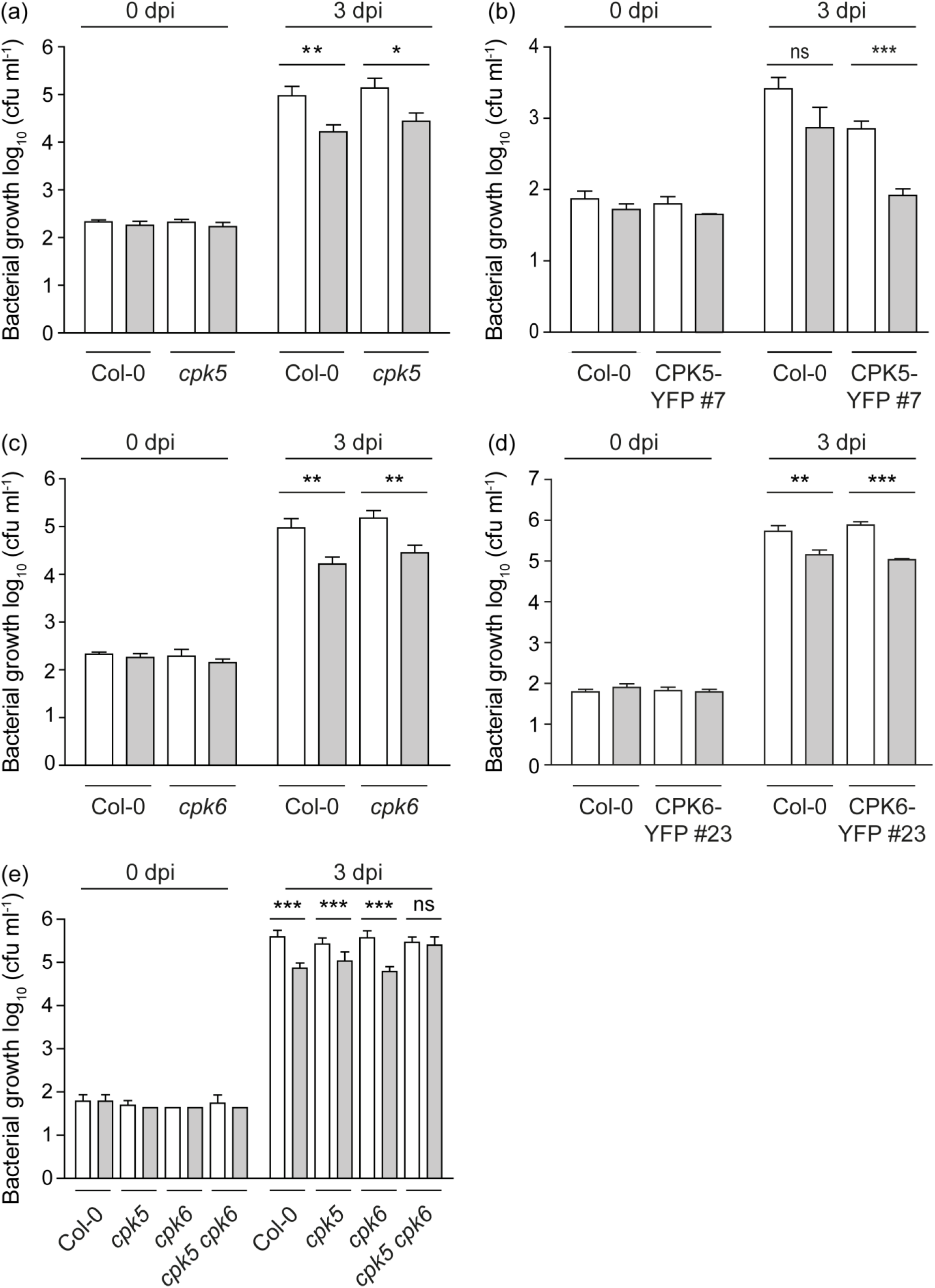
Enhanced CPK5-signaling triggers immune priming. (**a**-**e**) Six-week-old plants of Col-0, *cpk5* (**a**), *CPK5-YFP#7* (**b**), *cpk6* (**c**), *CPK6-YFP#23* (**d**), and *cpk5 cpk6* (**e**) were subjected to a priming infection by avirulent bacterial strain *Pseudomonas syringae pv. maculicola* ES 4326 avrRpm1 (*Psm avrRpm1*) (grey bars) or a mock control (10 mM MgCl_2_) (white bars) in local leaves. After two days, distal leaves were subjected to triggering infection with virulent strain *Psm* ES 4326. Bacterial numbers of the triggering strain were monitored 0 dpi and 3 dpi. Bars represent mean value ± SEM of 12 biological replicates. The asterisks indicate statistically significant differences in comparison to mock at 3 dpi (student’s t-test; ***, *P*<0.001; **, *P*<0.01; *, *P*<0.05; ns, not significant). The experiments were repeated three times with similar results.

**Fig. 4:**
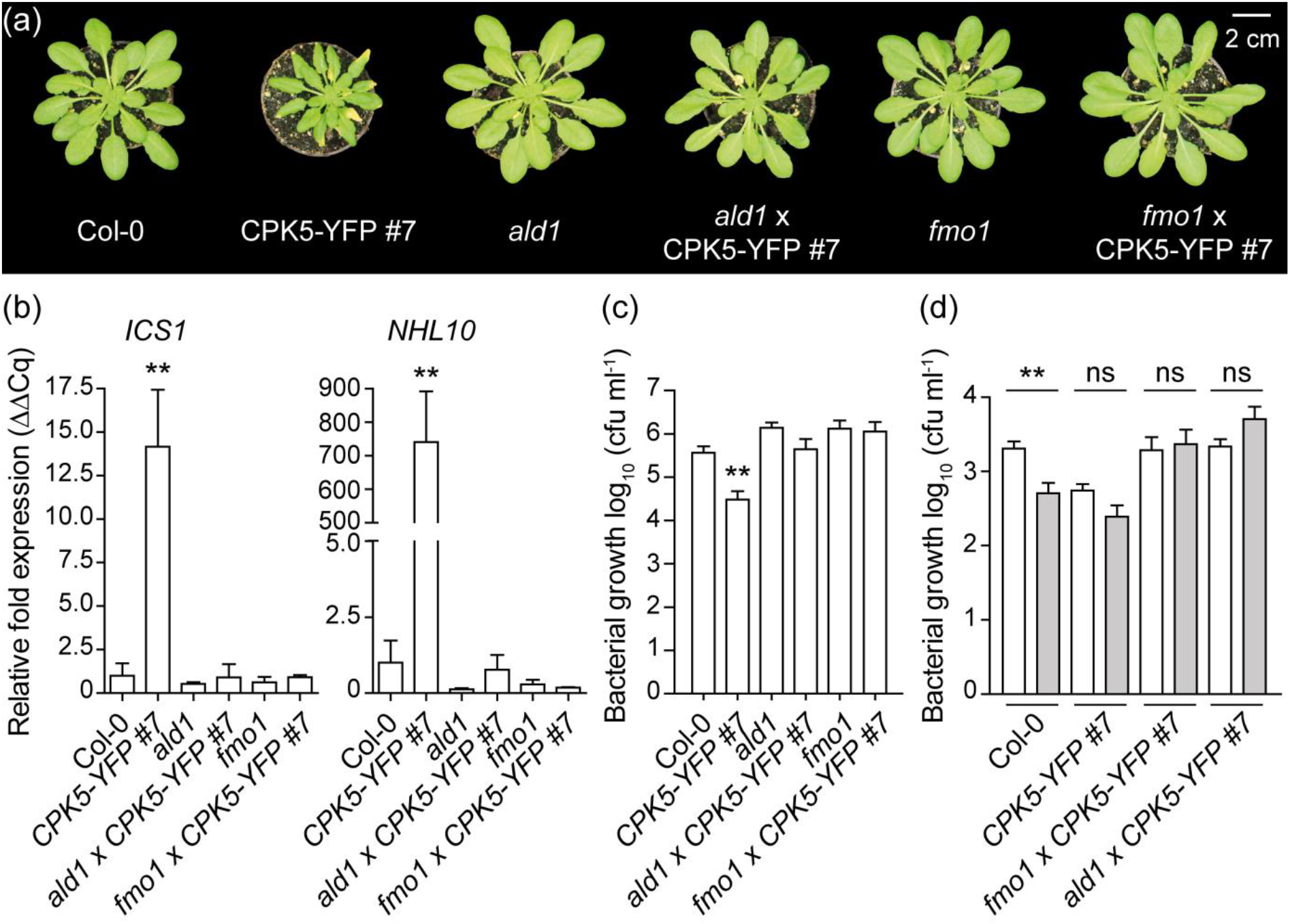
CPK5-mediated resistance is dependent on *ALD1* and *FMO1*. (**a**) Six-week-old plants of Col-0, *CPK5-YFP#7*, and derived crosses with *ald1* and *fmo1*. Scale = 2 cm. (**b**) Basal *ICS1* and *NHL10* gene expression of plants as shown in (a) was analyzed by RT-qPCR. Bars represent mean value ± SEM of three biological replicates. The asterisks indicate statistically significant differences in comparison to Col-0 (one-way ANOVA; Dunnett posttest; **, *P*<0.01). (**c**) Six-week-old plants as shown in (a) were inoculated with *Pseudomonas syringae pv. tomato* (*Pst*) DC3000. Bacterial numbers were monitored 0 dpi (data not shown) and 3 dpi. Bars represent mean value ± SEM of 12 biological replicates. The asterisks indicate statistically significant differences in comparison to Col-0 (one-way ANOVA; Dunnett posttest; **, *P*<0.01). The experiment was repeated three times with similar results. (**d**) Six-week-old plants as shown in (a) were subjected to priming infection with avirulent bacterial strain *Psm avrRpm1* (*Pseudomonas syringae pv. maculicola* (*Psm*) ES 4326 avrRpm1) or mock control (10 mM MgCl_2_) in local leaves. After two days, distal leaves were subjected to triggering infection with virulent strain *Psm* ES 4326. Bacterial numbers of the triggering strain were monitored 0 dpi (data not shown) and 3 dpi. Bars represent mean value ± SEM of 12 biological replicates. The asterisks indicate statistically significant differences in comparison to mock at 3 dpi (student’s t-test; **, *P*<0.01; ns, not significant). The experiment was repeated with similar results.

### CPK5-mediated priming is dependent on *ALD1/FMO1*-signaling

We next generated crosses between *ald1* and *fmo1* mutant lines, respectively, and *CPK5-YFP#7* and double homozygous plants were selected. Both crosses, *ald1* x *CPK5-YFP#7* and *fmo1* x *CPK5-YFP#7*, no longer show the reduced growth phenotype manifested in a smaller rosette, disordered leaf shape and lesion development caused by CPK5-YFP overexpression (Fig. **4a**). Also, basal expression of marker genes *ICS1* and *NHL10*, both elevated in *CPK5-YFP#7*, are reverted to Col-0 wild type level in the crosses (Fig. **4b**).

In standard bacterial growth assays using *Pst* DC3000 both single mutant lines, *ald1* and *fmo1,* are more susceptible to bacterial pathogens compared to wild type. In crosses, the enhanced resistance of *CPK5-YFP#7* is lost. Thus bacterial growth in *ald1* x *CPK5-YFP#7* is reverted to Col-0 wild type level and in *fmo1* x *CPK5-YFP#7* bacterial growth resembles that of the *fmo1* single mutant (Fig. **4c**). These data demonstrate that full CPK5-mediated resistance depends on *ALD1* and *FMO1*.

To address the dependency of CPK5-mediated SAR on *ALD1* and *FMO1*, priming experiments were conducted in temporal and special resolution as described above. Priming as observed in Col-0 and the ‘super-priming’ phenotype of *CPK5-YFP#7* was entirely absent in *ald1* x *CPK5-YFP#7* and *fmo1* x *CPK5-YFP#7* (Fig. **4d**). Instead, inoculated plants showed increased susceptibility like un-primed Col-0 or became even more susceptible (*fmo1* x *CPK5-YFP#7*), consistent with FMO1 enzyme catalyzing the reversion of SAR marker metabolite Pip to NHP during SAR (Hartmann *et al.*, 2018).

### CPK5-mediated priming depends on *SARD1*

We next assessed whether CPK5-mediated ‘super-priming’ requires SAR key transcription factor *SARD1*. Our expression analysis revealed a high constitutive level of *SARD1* transcript in line *CPK5-YFP#7* compared to Col-0. *SARD1* transcript accumulation is absent in priming-deficient crosses *ald1* x *CPK5-YFP#7* and *fmo1* x *CPK5-YFP#7* and also in the *ald1* and *fmo1* single mutants whereas basal *SARD1* expression is slightly reduced in *cpk5* compared to Col-0 (Fig. **5a**).

**Fig. 5:**
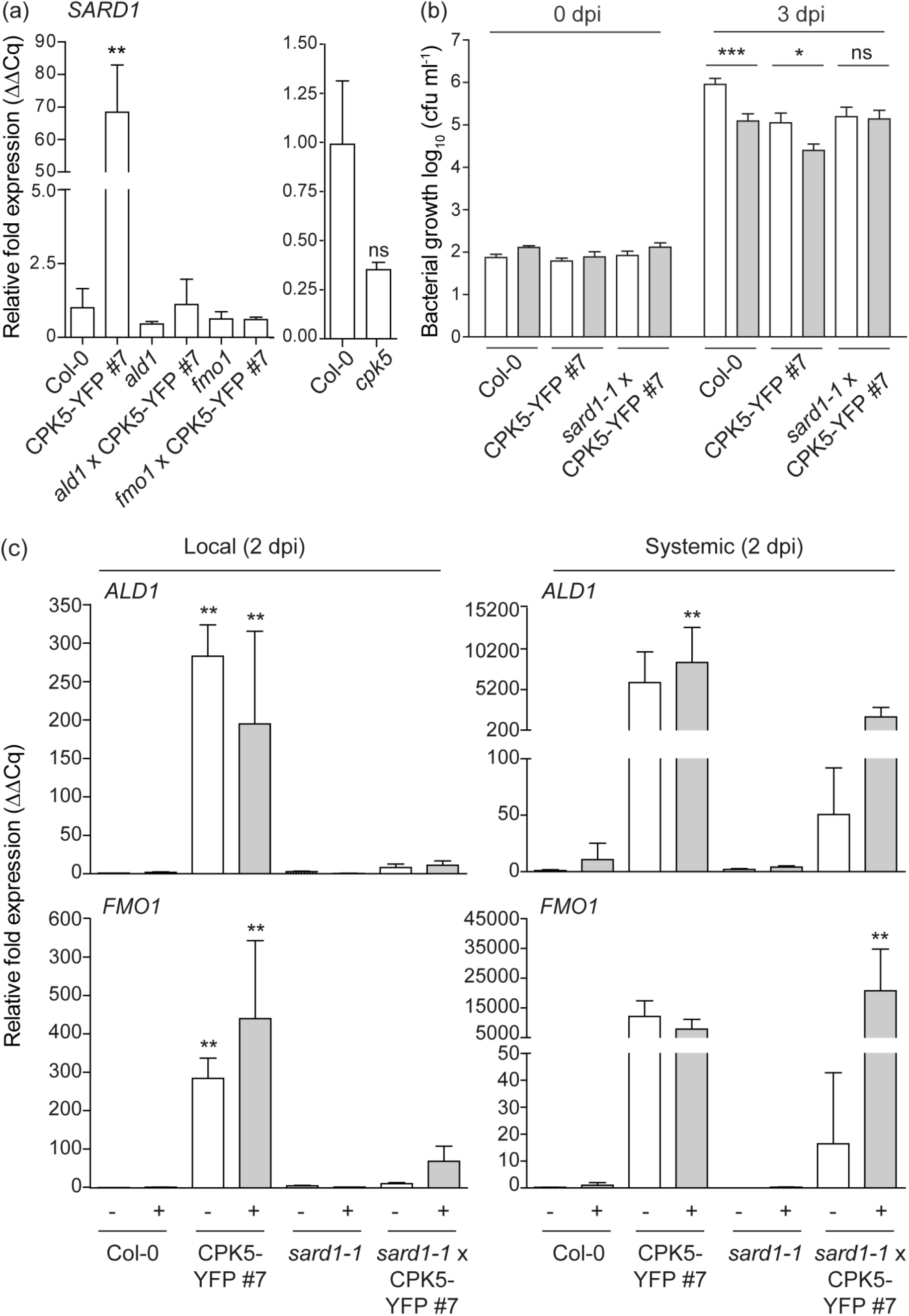
CPK5-dependent ‘super-priming’ requires *SARD1*. (**a**) Basal *SARD1* gene expression in six-week-old plants of Col-0, *CPK5-YFP#7*, and derived crosses with *ald1* and *fmo1* (left panel) and *cpk5* (right panel), respectively, were analyzed by RT-qPCR. Bars represent mean value ± SEM of three biological replicates. The asterisks indicate statistically significant differences in comparison to Col-0 (one-way ANOVA; Dunnett posttest; **, *P*<0.01; ns, not significant). (**b**) Six-week-old plants of Col-0, *CPK5-YFP#7*, and *sard1*-1 x *CPK5-YFP#7* were subjected to priming and triggering infection as described in (Fig. 3 (b)). Bacterial numbers were monitored 0 dpi (data not shown) and 3 dpi of the triggering strain. Bars represent mean value ± SEM of 12 biological replicates. The asterisks indicate statistically significant differences in comparison to mock at 3 dpi (student’s t-test; **, *P*<0.01; *, *P*<0.05; ns, not significant). The experiment was repeated with similar results. (**c**) Local and systemic *ALD1* and *FMO1* expression of Col-0, CPK5-YFP#7, *sard1-1* and *sard1-1* x CPK5-YFP#7 was analyzed by qRT-PCR 2 dpi after mock (white bars) or flg22 (grey bars) stimulus as described in Fig. 2 (a). Bars represent mean value ± SEM of three biological replicates. The asterisks indicate statistically significant differences in comparison to Col-0 mock (one-way ANOVA; Dunnett posttest; **, *P*<0.01). The experiment was repeated with similar results.

To investigate whether *SARD1* is required for CPK5-mediated priming or ‘super-priming’, a cross between *sard1-1* and *CPK5-YFP#7* was generated and double homozygous plants were analyzed. In the crossed lines, local and systemic *ALD1* and *FMO1* gene expression at 2 dpi, which is constitutively elevated in *CPK5-YFP#7*, is entirely absent (Fig. **5c**). The same is observed for *PR1* and *FRK1* (Fig. **1b**). Remarkably, in *sard1*-*1* x *CPK5-YFP#7* the systemic *ALD1* and *FMO1* expression is low in unstimulated conditions (resembling Col-0), but upon flg22-stimulation both gene transcripts accumulate to high levels comparable to those of the CPK5-overexpressing line. These data correlate with the phenotype of *sard1*-*1* x *CPK5-YFP#7* plants, which display a reduced rosette diameter and lesion development compared to *CPK5-YFP#7* (Fig. **1a**), and which also show enhanced resistance to the virulent bacterial pathogen *Pst* DC3000 (Fig. **1c**).

Priming experiments for bacterial growth in SAR reveal that *sard1*-*1* x *CPK5-YFP#7* plants are still able to repress bacterial growth to the level of the CPK5-overexpression line. However, in the absence of *SARD1*, these plants are unable to induce ‘super-priming’ observed in the CPK5-overexpressing line (Fig. **5b**). Thus, ectopic activation of CPK5 throughout the entire plant induces immunity against bacterial pathogens. Although defense is already activated by CPK5-YFP in distal leaves, an additional SARD1-dependent signal is relayed. These data suggest that CPK5 signaling mediates both, rapid local and long-term distal resistance responses, to restrict pathogen growth. *SARD1*, a transcriptional regulator whose gene itself is induced late during pathogen infection, is required only for the latter. Whereas in *sard1*-*1* x *CPK5-YFP#7* CPK5-mediated enhanced basal resistance is unaltered, plants miss the late additional resistance benefit of having been primed.

### CPK5 is a highly responsive calcium-activated enzyme

The calcium concentration at which a CDPK displays half-maximal phosphorylating kinase activity, K50 [Ca^2+^], reflects an isoform-specific biochemical parameter and differs even between close homologs within the CDPK gene family. A low K50 for calcium is indicative for an enzyme that has adopted its catalytically active conformation at low concentrations of calcium. In standard *in vitro* kinase assays with recombinant enzymes toward synthetic peptide substrate Syntide 2 the K50 [Ca^2+^] for CPK5 is as low as ∼94 nM and for CPK6 ∼186 nM (Fig. **6a,c**). The intracellular calcium concentration in unstimulated plants is reported to ∼100 nM (Stael *et al.*, 2012; Costa *et al.*, 2018). Thus, at the resting cytoplasmic calcium concentration CPK5 but not yet CPK6 activity is already highly responsive to subtle concentration changes. A likewise low K50 [Ca^2+^] is observed with peptide substrate encompassing S148 from the CPK5 *in vivo* phosphorylation target RBOHD (Dubiella *et al.*, 2013; Kadota *et al.*, 2014) (Fig. **6b,d**). Likewise, similar low K50 [Ca^2+^] is determined for fusion protein CPK5-YFP for both substrates (Fig. **S5**).

**Fig. 6:**
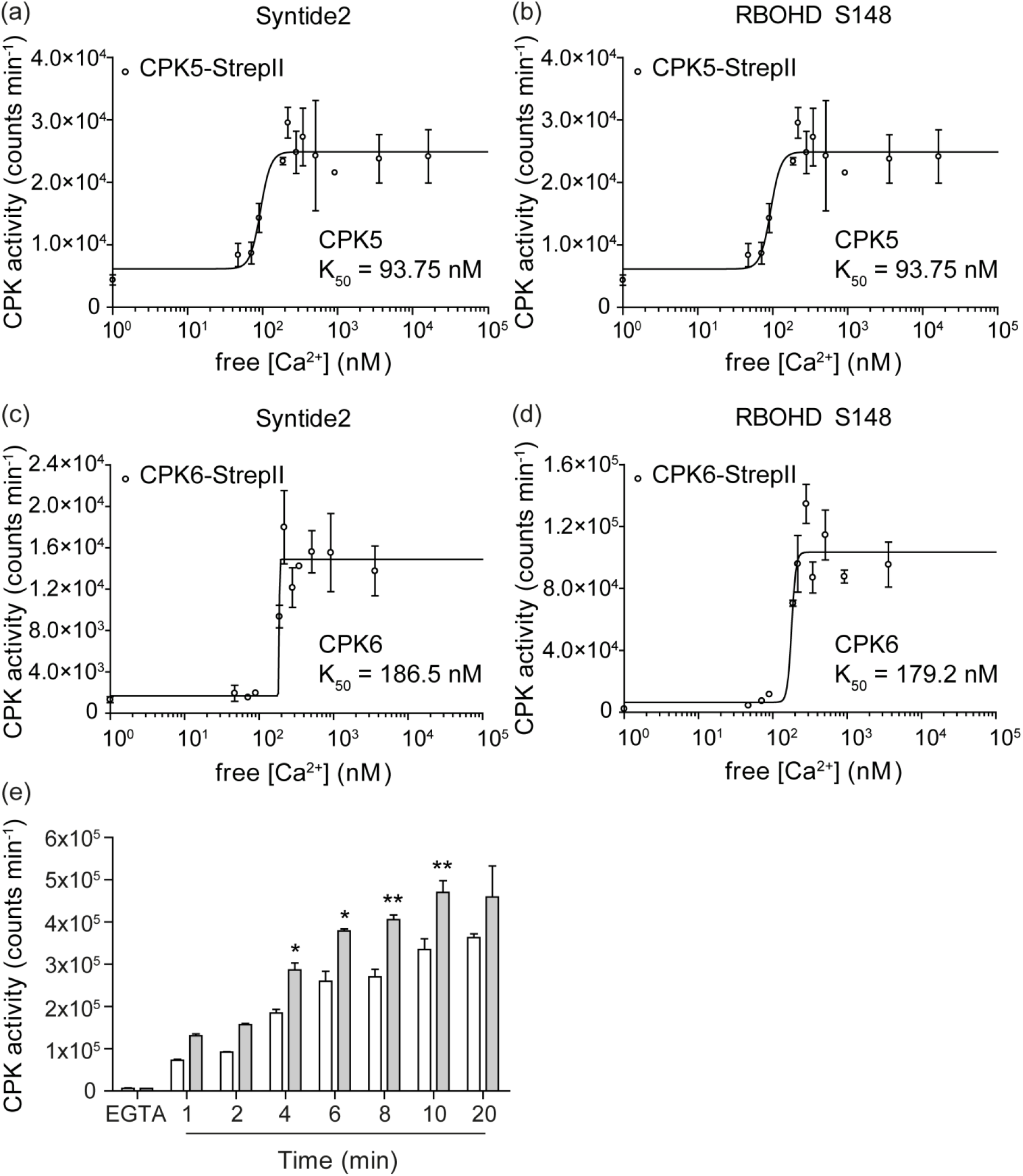
CPK5 displays high kinase activity at low Ca^2+^-concentrations. (**a**), (**b**) Ca^2+^-dependent protein kinase activity of CPK5-StrepII. Kinase activity of affinity-purified recombinant protein CPK5-StrepII (25 nM) was analyzed in an *in vitro* kinase assay toward 10 µM substrate peptide Syntide 2 (a) and RBOHD S148 (10 aa peptide encompassing S148) (b), respectively, for 20 min in a series of increasing Ca^2+^ concentrations as indicated and K50 was determined. Data represent mean value ± SEM of two technical replicates. The experiment was repeated with similar results. (**c**), (**d**) Ca^2+^-dependent protein kinase activity of CPK6-StrepII. Kinase activity of affinity-purified recombinant protein CPK6-StrepII (25 nM) was analyzed in an *in vitro* kinase assay toward Syntide 2 (c) and RBOHD S148 (d), respectively, as described for (a), (b). (**e**) Ca^2+^-dependent protein kinase activity of affinity-purified recombinant CPK5-StrepII or CPK5-YFP-StrepII (25 nM protein) was assessed over a fast time kinetic from 0 to 20 min in an *in vitro* kinase assay with 10 µM substrate peptide RBOHD S148 without calcium or at a fixed saturating Ca^2+^ concentration of 0.5 µM. Bars represent mean value ± SEM of two technical replicates. The asterisks indicate statistically significant differences of CPK5-YFP-StrepII in comparison to CPK5-StrepII (two-way ANOVA; Bonferroni posttest; *, *P*<0.05; **, *P*<0.01). The experiment was repeated with similar results.

Remarkably, the CPK5-YFP fusion protein displays higher kinase activity at low [Ca^2+^] concentrations compared to CPK5 in a rapid kinetic assay (Fig. **6e**). This is consistent with an interpretation that CPK5-YFP undergoes more rapidly (and at lower cytoplasmic calcium concentrations) a calcium-induced conformational change to adapt and stabilize the active conformation compared to the native enzyme. In accordance with this analysis CPK5-YFP transgenic plants display constitutive biochemical (in-gel kinase) activity, and a further increase in kinase activity was induced upon flg22 treatment of these plants (Dubiella *et al.*, 2013).

## Discussion

Systemic acquired resistance represents a status of preparedness of the entire plant foliage to a broad spectrum of microbial pathogens induced by a preceding infection at a local site. This preparedness is corroborated by more rapid and vigorous activation of defense responses. Because long-term defense activation may come at the cost of growth retardation and a delay of plant development, the decision to switch in the SAR mode has to be tightly controlled. SA plays a pivotal role in plant immunity, and SA accumulation is undoubtedly a key executor to establish and maintain SAR. However, SA is not considered to be the one and only signal molecule to be transported through the plant. Thus, the switch to SAR requires an information relay of the ‘having been attacked’ perception from local to distal plant sites as well as the initiation and maintenance of an SA-accumulating defense loop that manifests a ‘being prepared when it happens again’ information (Hake & Romeis, 2018).

### CPK5 signaling triggers *SARD1*-dependent SAR and mediates ‘super-priming’

Plants overexpressing CPK5-YFP show increased basal resistance to the virulent bacterial pathogen *Pst*. CPK5 signaling-mediated immunity is thereby dependent on SA. In crosses of the *CPK5-YFP#7* line to mutants *ics1, eds5, pad4*, and *npr1-1* implicated in SA biosynthesis and signaling in PTI and ETI, CPK5-dependent defense marker gene expression and resistance to virulent *Pst* is compromised and plant growth is reverted to wild type (Fig. **1**). CPK5 can, therefore, be placed upstream of PAD4 and the cascade of SA-dependent defense reactions.

Here we show that CPK5-YFP overexpressing plants constitutively express SAR marker genes *ALD1* and *FMO1* at late time points in distal leaves, and these plants also accumulate Pip (Fig. **2**, Fig. **S2**). Plants of enhanced CPK5-signaling contain therefore all essential components for a Pip- and SA-dependent signaling amplification loop via *ALD1*/*FMO1*/*ICS1*. Consistently, CPK5-YFP overexpressing lines show a ‘super-priming’ phenotype of enhanced bacterial resistance in the context of SAR (Fig. **3**, **4**, **5**). Priming and ‘super-priming’ is entirely lost in crosses *ald1* x *CPK5-YFP#7* and *fmo1* x *CPK5-YFP#7* (Fig. 4d), and the benefit of enhanced CPK5-signaling is lost in both, basal and systemic resistance (Fig. **4c,d**).

Additionally, enhanced CPK5-signaling results in a constitutive accumulation of *SARD1* transcript, validating the constitutive high *ALD1* and *FMO1* expression levels in these lines even in absence of any pathogen-related stimulation. Comparably low systemic *SARD1* expression and low *ALD1* and *FMO1* transcript levels are detected in *cpk5* (Fig. **2b**). Plants that overexpress *SARD1* are reported to be more resistant to bacterial infection in basal immunity and to accumulate SA (Zhang *et al.*, 2010), similar to what is observed for CPK5-overexpressing lines. Interestingly, high constitutive transcript levels of *ALD1* and *FMO1* in *CPK5-YFP#7* are reverted in a *sard1* background (Fig. **5c**), mimicking the *sard1* single mutant (Sun *et al.*, 2015). Remarkably, upon an flg22 immune stimulus, systemic *ALD1* and *FMO1* expression at 2 dpi is induced to a comparably high level as seen for enhanced CPK5-signaling in *CPK5-YFP#7* (Fig. **5c**). These data suggest that a local pathogen attack induces basal resistance which includes CPK5, CPK6, and in case other CPKs. In addition, CPK5 participates in signal propagation and in the control of a distal switch into SAR. Yet, in the absence of *SARD1*, SAR cannot be maintained.

This interpretation is mirrored by SAR bacterial growth phenotypes of the respective crossing lines. Plants of *ald1* x *CPK5-YFP#7* and *fmo1* x *CPK5-YFP#7* lost both, the CPK5-mediated enhanced basal resistance and systemic priming (Fig. **4c,d**). In *sard1* x *CPK5-YFP#7* the enhanced CPK5-dependent basal resistance from *CPK5-YFP#7* is retained, but a memory of ‘having been attacked’, the *SARD1*-dependent additional (super-) priming which is corroborated by a constitutive *ALD1* and *FMO1* expression, is lost (Fig. **5b**). These data link CPK5-signaling with SARD1 function, and both are required to mount and maintain the primed plant state manifesting the immune memory. *sard1* plants still respond to flg22 with local and systemic immune reactions and thus are able to synthesize and propagate an immune signal. However, the ultimate switch into SAR through a SA- and Pip-dependent self-containing amplification loop of *ALD1*/*FMO1*/*ICS1* cannot be accomplished in the absence of *SARD1*.

SARD1 is a key regulatory transcription factor that binds to the promoters of a large number of genes, for which a positive or negative role in systemic plant resistance can be attributed, including *ICS1*. Its expression is tightly controlled by positive and negative regulation in a temporally and spatially manner and occurs late in systemic tissue (Wang *et al.*, 2011; Zheng *et al.*, 2015; Sun *et al.*, 2018; Zhou *et al.*, 2018). It is unclear yet, whether *SARD1* expression is also controlled by post-transcriptional mechanisms that may directly impact *SARD1*-dependent SAR, for example by phosphorylation through CPK5.

### CPK5 links the calcium-regulatory network with immune signaling

The role of calcium-signaling, well characterized during local PAMP-induced immune reactions, is less clear in late systemic signaling and the switch to SAR. Regulatory calcium-binding proteins such as CMLs and AGP5, in particular, those that are under transcriptional control of *SARD1/CBP60g* (Truman & Glazebrook, 2012; Aldon *et al.*, 2018), may depend on temporal and spatial distinct intracellular calcium conditions from those of initiating local calcium burst. Interestingly, calmodulin-dependent transcriptional regulators CAMTA3 and CBP60a have been described as negative regulators in the control of long-term transcriptional reprogramming of defense genes (Galon *et al.*, 2008; Truman & Glazebrook, 2012; Sun *et al.*, 2015). These data implicate that the intracellular calcium status is essential also in the control of SAR.

CPK5 and CPK6 classify to the CPDK subfamily I and comply with a consensus CDPK enzyme with 4 canonical EF-hand motifs (Cheng *et al.*, 2002). In biochemical assays the catalytic activities of CPK5 and CPK6 are calcium-dependent. A remarkably low K50 [Ca^2+^] of 93 nM was determined for CPK5 and of 186 nM for CPK6 (Fig. **6a,c**). These data indicate that in particular CPK5 is most sensitive to subtle [Ca^2+^] changes around the intracellular resting calcium concentration. Thus, this low K50 [Ca^2+^] may explain why CPK5 (but not CPK6) is part of a signal propagation mechanism from local to distal sites via a CPK5/RBOHD-driven auto-activation circuit (Dubiella *et al.*, 2013), why overexpression of CPK5 (but not of CPK6) induces SAR, and why CPK5 (but not CPK4, CPK6, and CPK11) is required for defense responses in autoimmune mutants such as in *exo70B1* (Liu *et al.*, 2017). Thus, CPK6 (here: K50 [Ca^2+^] of 186 nM) and other CPKs such as CPK4 and CPK11 (reported K50 [Ca^2+^] of ∼ 3 µM and ∼ 4 µM, respectively (Boudsocq *et al.*, 2012)) may become fully activated and contribute to defense activation upon an intracellular calcium burst for example as consequence of a direct local pathogen attack or exposure to PAMP flg22. But these enzymes may not be suited to decode subtle calcium changes when propagating a signal or activating SAR in distal tissue. CPK6, the phylogenetically closest homolog to CPK5, has predominantly been implicated in guard cell function during the control of the stomatal aperture (Mori *et al.*, 2006; Brandt *et al.*, 2012; Ye *et al.*, 2013). The CPK6 overexpressing line *CPK6-YFP#23* does neither show constitutive SAR marker gene expression nor enhanced SAR towards bacterial pathogens (Fig. **2a**, **3d**). Consistently, ROS-generation driving immune signal propagation has been shown to be compromised in *cpk5* but not in *cpk6* (Dubiella *et al.*, 2013).

### CPK5 protein amount and catalytic activity contribute to the manifestation of priming

The underlying mechanism that manifests SAR is a matter of ongoing research and the accumulation and activation of signaling molecules, the modifications at the chromatin level and at the transcriptome may all contribute to immune memory (Conrath *et al.*, 2015; Hilker *et al.*, 2015; Martinez-Medina *et al.*, 2016; Reimer-Michalski & Conrath, 2016; Hake & Romeis, 2018).

A role of mitogen-activated protein (MAP) kinases MPK3 and MPK6 has been demonstrated in chemically-induced resistance. Priming correlated with enhanced *MPK* transcript levels and the accumulation of (yet inactive) MPK3 and MPK6 proteins. Exposure to a triggering stimulus not only led to enhanced MPK biochemical activation but also to increased MPK-dependent downstream signaling, resulting in more prominent defense reactions. Consistently, priming responses were compromised or lost in mutant lines of *mpk3* and *mpk6* (Beckers *et al.*, 2009). MPK3 and MPK6 were shown to contribute to SAR via an *ALD1*-pipecolic acid regulatory loop (Y. Wang *et al.*, 2018). Furthermore, an interplay between MPK and CPK signaling has been demonstrated for innate immune signaling in PTI (Boudsocq *et al.*, 2010), where exemplary defense gene activation was shown to be either MPK-specific, or CPK-specific, or to be controlled through joint MPK- and CPK-signaling. CPK5 signaling contributed to the activation of transcriptional regulators WRKY8, 28, and 48 upstream of WRKY46 and CPK5 phosphorylates recombinant WRKY28 and WRKY48 proteins (Gao *et al.*, 2013). Thus, it is conceivable that transcriptional regulators downstream of protein kinase signaling become increasingly essential in the transition from ETI to SAR. Interestingly, Chip-Seq analysis revealed promoter sequences recognized by SARD1 in primed plants. Among SARD1-binding sequences have been identified promoters of signaling genes *MPK3* and *CPK4*, in addition to promoters of genes of the SA/Pip amplification loop (*ALD1*/*FMO1*/*ICS1*), and of promoters of genes mediating general SA-dependent PTI and ETI defense signaling (*EDS1, PAD4, NPR1)* (Sun *et al.*, 2015). These data corroborate that the accumulation of (potentially not yet fully active) signaling proteins such as MPKs and CPKs are part of the repertoire to manifest a primed plant status.

In the CPK5-overexpressing line *CPK5-YFP#7* the enzyme accumulates to a high protein amount that is even stronger in line CPK5-YFP # 2 (Dubiella *et al.*, 2013). Additionally, the CPK5-YFP fusion protein is biochemically more active at a given low cytoplasmic [Ca^2+^] level than native CPK5, which correlates with an observed constitutive biochemical phosphorylation activity of CPK5-YFP in these lines (Dubiella *et al.*, 2013). Taken together, our data are consistent with the interpretation that CPK5-YFP mimics a pre-formed (‘primed’), phosphorylation competent state (synonymous to a so-called ‘protein mark’), which in dependency of *SARD1* is responsible for ‘super-priming’.

Our data are in accordance with a model where CPK5 guards the cytoplasmic calcium status in resting cells facilitated by its low K50 [Ca^2+^] (Fig. **7**). Following a primary pathogen attack and subsequent rise in cytoplasmic [Ca^2+^], CPK5 predominantly acquires its open conformation, and local defenses and basal immunity are established, in case in joint function with other locally activated CPKs. In ETI, the information of ‘having been attacked’ is spread via a CPK5-mediated calcium/ROS propagation mechanism and meets the command of ‘having to defend’ mediated through the synthesis and accumulation of SA and Pip / NHP defense metabolites, which in the presence of SARD1 lead to a switch into and maintenance of SAR in systemic plant tissue. It is conceivable that during the immune memory CPK5 adopts a primed enzyme state, likely manifested through a distinct intramolecular pattern of protein phosphorylation combined with a change in protein conformation that allows a more rapid enzyme transition to the fully active state upon a triggering stimulus, for example during a secondary pathogen attack (Fig. **7**).

**Fig. 7:**
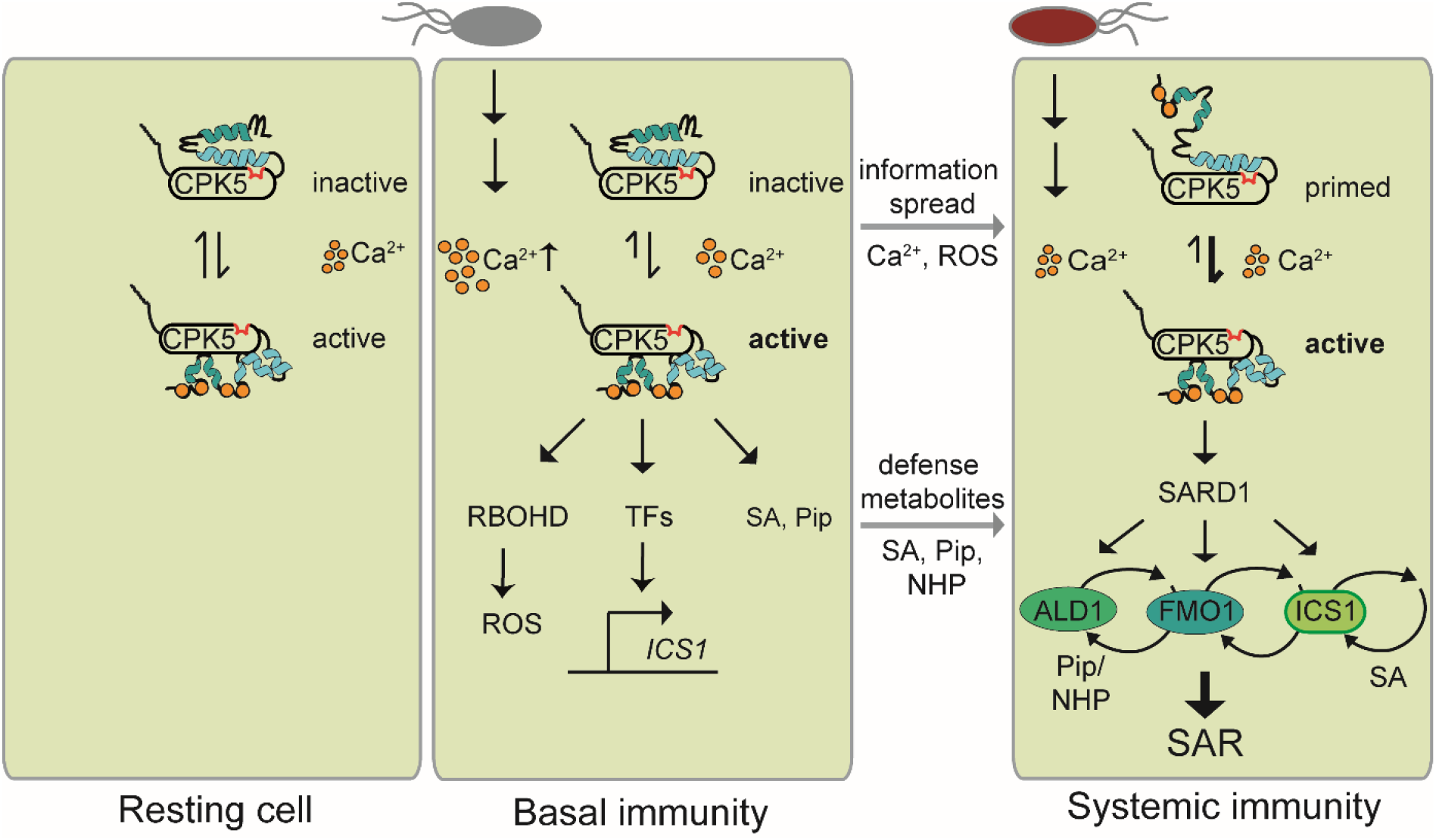
Scheme of Ca^2+^-dependent CPK5 activation and function in immune responses, priming, and SAR.

Thus, CPK5-signaling, already required for the spatial defense signal spread into the entire foliage of a plant, may also contribute to the temporal switch - via the linkage of calcium signaling to the activation of *SARD1* – to induce (reversible) SAR. Whether *SARD1* is solely activated on the transcriptional level or also on the post-translational level, e.g. upon direct interaction and phosphorylation of SARD1 protein through CPK5, remains to be shown.

## Acknowledgments

This research was funded by Deutsche Forschungsgemeinschaft (DFG) within Collaborative Research Centre SFB973 to T.R. We thank Jennifer Bortlik for the transformation of *Arabidopsis* with the CPK6-YFP construct and Katharina Hake for critical reading and discussion of the manuscript. *Psm* and *Psm avrRpm1* strains were kindly provided by Jürgen Zeier (University Düsseldorf), the *cpk5 cpk6* double mutant line was kindly provided by Marie Boudsocq (Institute of Plant Sciences Paris Saclay).

## Author contributions

T.G., T.R. conceived and designed the research. T.G., S.S., F-P.S., B.C. performed experiments. T.G., B.C., F-P.S., T.R. analyzed data. T.G. and T.R. wrote the manuscript.

---

**Fig. S1:**
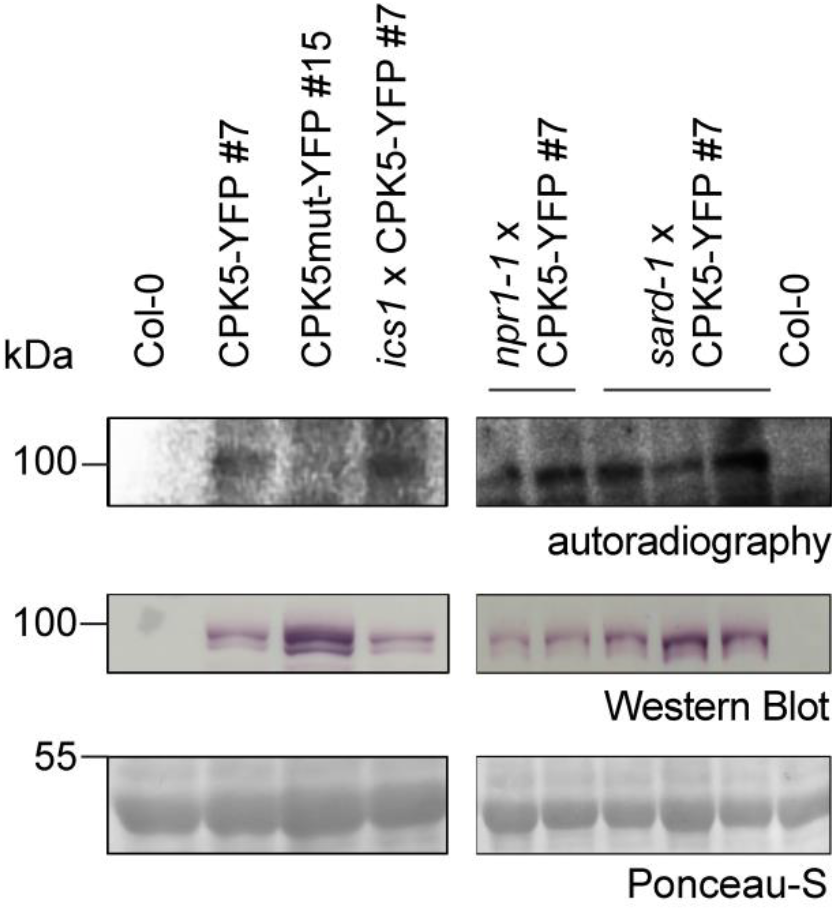
CPK5-YFP is expressed and retains constitutive protein kinase activity in crossed lines. Total protein of two-week-old seedlings of Col-0, CPK5-YFP #7, CPK5mut-YFP # 15, and derived *Arabidopsis* crossing lines as indicated were separated by SDS-PAGE and subjected to an in-gel kinase assay using myelin basic proteins as a substrate or to immunoblotting. Proteins were visualized by autoradiography (upper panel) or analyzed by immunostaining with α-YFP (middle panel). The amount of protein was visualized by Ponceau-S staining (lower panel).

**Fig. S2:**
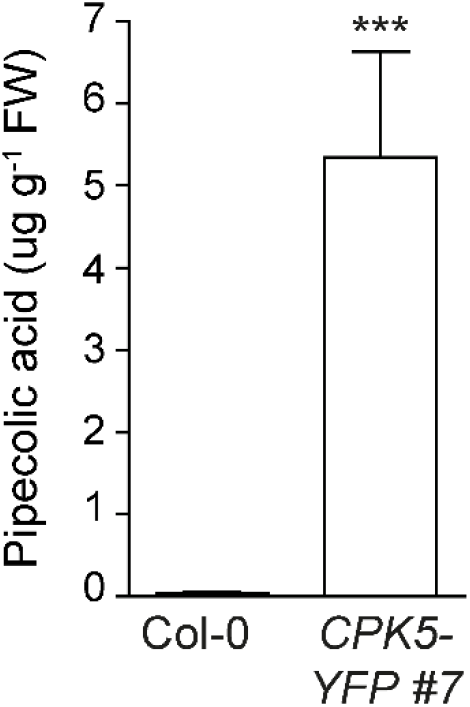
Overexpression of CPK5 results in the accumulation of pipecolic acid. Eight pools of five plants of Col-0 and *CPK5-YFP* #7 were analyzed for their pipecolic acid contents using the EZ:faast protocol for extraction and GC-MS analysis. The asterisk indicates statistically significant differences in comparison to Col-0 (student’s t-test, *** *P*<0.001).

**Fig. S3:**
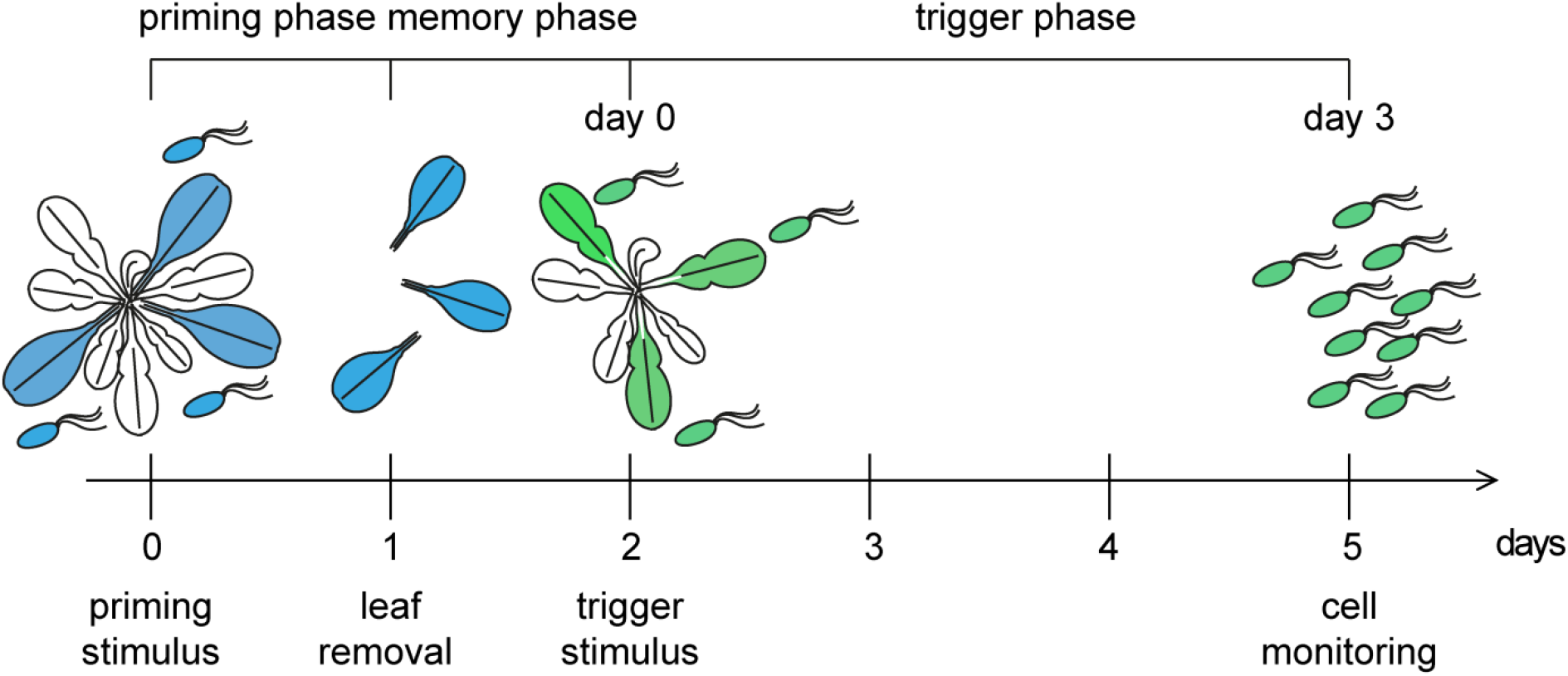
Experimental outline to assess temporal and spatial distinct responses required for priming of SAR. Plants are subjected to a primary stimulation (*priming stimulus*) by treatment with 200 nM flg22 or infection with avirulent *Psm* ES 4326 *avrRpm1* and a control (infiltration of 10 mM MgCl_2_ as mock treatment) on three fully developed lower leaves (shown in blue). After 24 h these leaves were removed. After two days, three subsequent systemic leaves (shown in green) were subjected to a secondary stimulation (*triggering stimulus*) by infection with virulent *Psm* ES 4326 and the corresponding equality of infiltration events was monitored (corresponds to 0 dpi of cell monitoring). After five days, comparing to 3 dpi after the triggering stimulation, bacterial growth of virulent Psm ES 4326 was monitored.

**Fig. S4:**
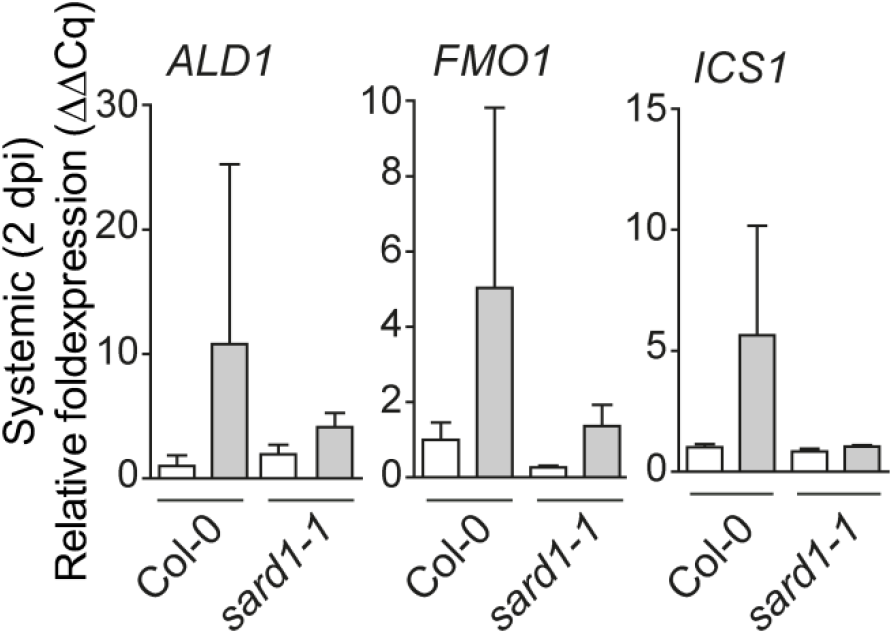
Systemic defense gene expression at two dpi is reduced in *sard1-1*. Three local leaves of six-week-old plants of Col-0 and *sard1-1* were infiltrated with 10 mM MgCl_2_ (mock/white bars) or 200 nM flg22 (grey bars). After two days, three systemic leaves were harvested and gene expression was analyzed by qRT-PCR. Bars represent mean value ± SEM of three biological replicates.

**Fig. S5:**
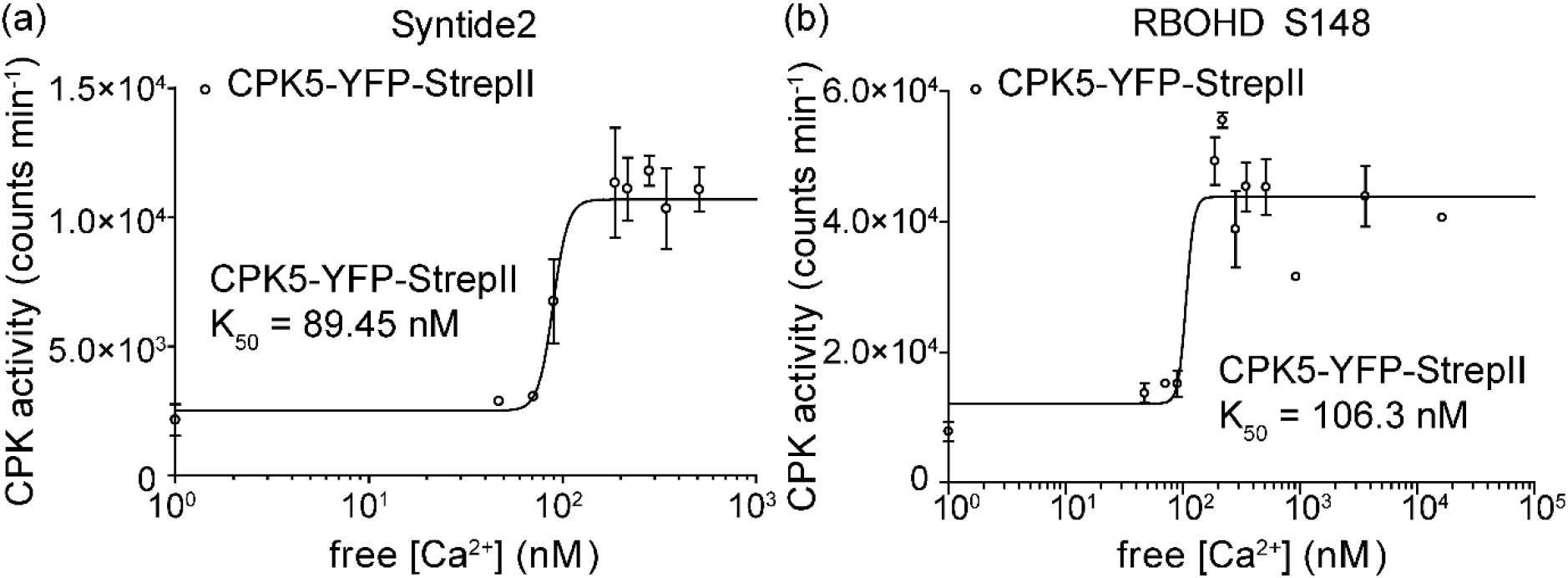
Ca^2+^-dependent protein kinase activity of CPK5-YFP-StrepII. Kinase activity of affinity-purified recombinant protein CPK5-YFP-StrepII (25 nM) was analyzed in an *in vitro* kinase assay toward 10 µM substrate peptide Syntide 2 (a) and RBOHD S148 (10 aa peptide encompassing S148) (b), respectively, for 20 min in a series of increasing Ca^2+^ concentrations as indicated and K50 was determined. Data represent mean value ± SEM of two technical replicates. The experiment was repeated with similar results.

**Table S1:**
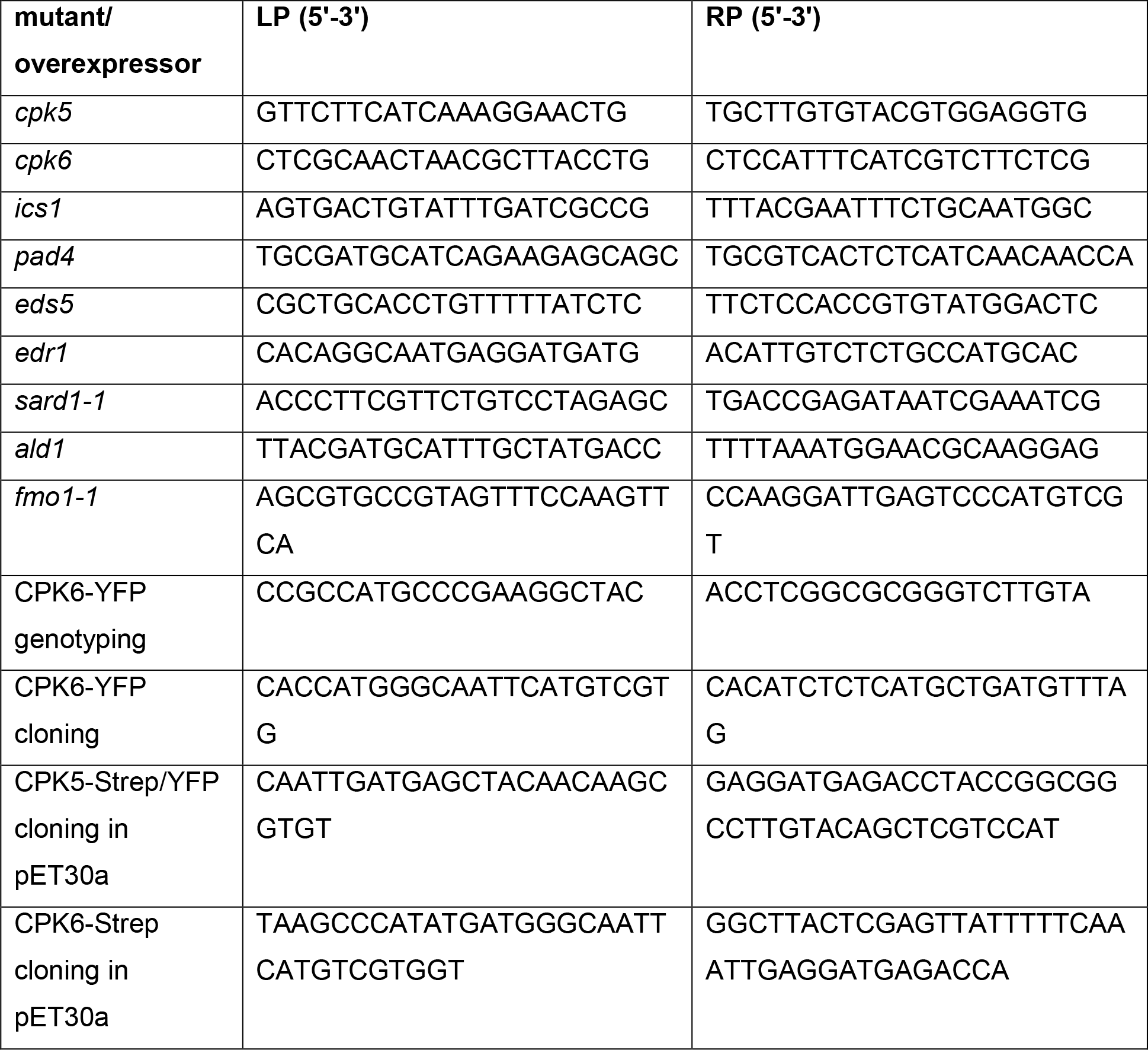
Primers used for genotyping of T-DNA insertion lines and cloning.

**Table S2:**
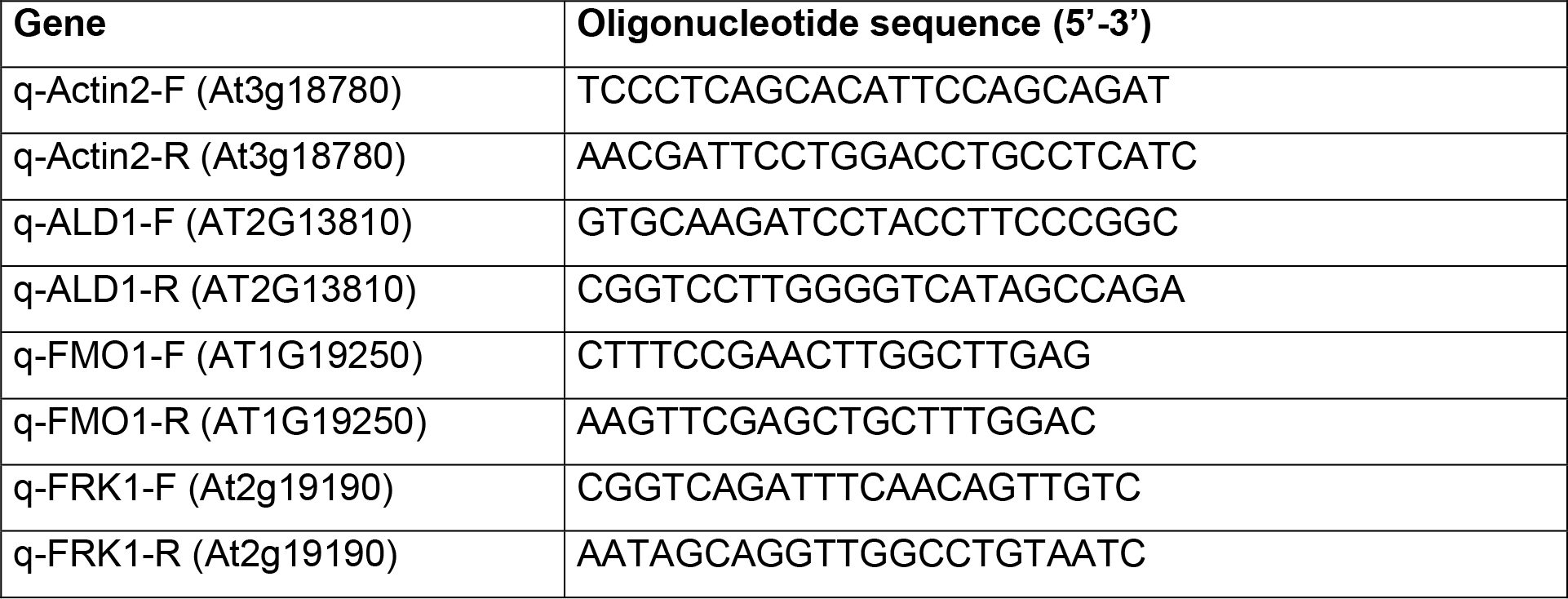
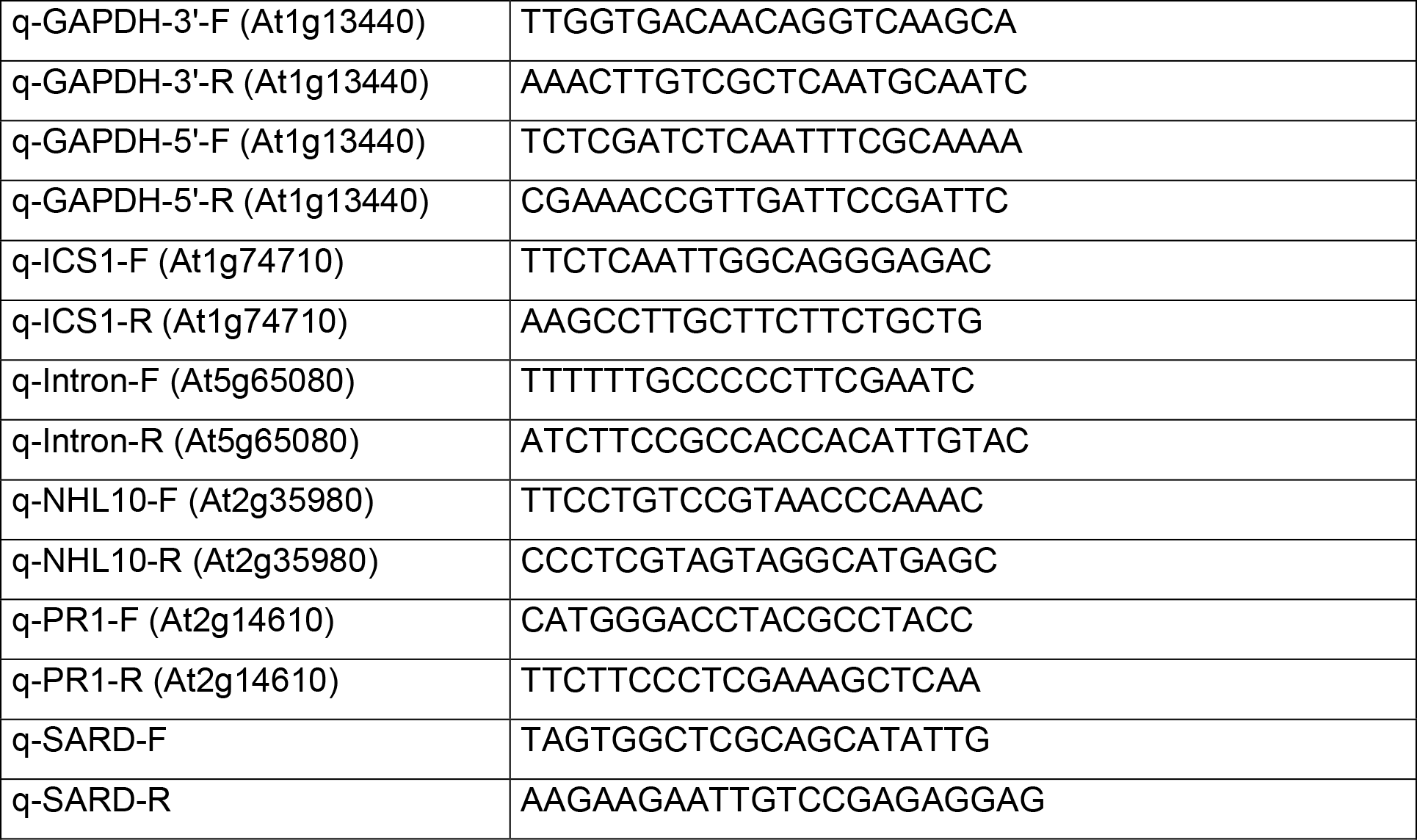
Primers used for RT-qPCR analysis.

## References

Aldon D, Mbengue M, Mazars C, Galaud JP. 2018. Calcium signalling in plant biotic interactions. Int J Mol Sci 19(3).

Beckers GJ, Jaskiewicz M, Liu Y, Underwood WR, He SY, Zhang S, Conrath U. 2009. Mitogen-activated protein kinases 3 and 6 are required for full priming of stress responses in arabidopsis thaliana. Plant Cell 21(3): 944–953.

Bernsdorff F, Doring AC, Gruner K, Schuck S, Brautigam A, Zeier J. 2016. Pipecolic acid orchestrates plant systemic acquired resistance and defense priming via salicylic acid-dependent and -independent pathways. Plant Cell 28(1): 102–129.

Boudsocq M, Droillard MJ, Regad L, Lauriere C. 2012. Characterization of arabidopsis calcium-dependent protein kinases: Activated or not by calcium? Biochem J 447(2): 291–299.

Boudsocq M, Willmann MR, McCormack M, Lee H, Shan L, He P, Bush J, Cheng SH, Sheen J. 2010. Differential innate immune signalling via ca^2+^ sensor protein kinases. Nature 464(7287): 418–422.

Brandt B, Brodsky DE, Xue S, Negi J, Iba K, Kangasjarvi J, Ghassemian M, Stephan AB, Hu H, Schroeder JI. 2012. Reconstitution of abscisic acid activation of slac1 anion channel by cpk6 and ost1 kinases and branched abi1 pp2c phosphatase action. Proc Natl Acad Sci U S A 109(26): 10593–10598.

Cheng SH, Willmann MR, Chen HC, Sheen J. 2002. Calcium signaling through protein kinases. The arabidopsis calcium-dependent protein kinase gene family. Plant Physiol 129(2): 469–485.

Conrath U. 2006. Systemic acquired resistance. Plant Signal Behav 1(4): 179–184.

Conrath U, Beckers GJ, Langenbach CJ, Jaskiewicz MR. 2015. Priming for enhanced defense. Annu Rev Phytopathol 53: 97–119.

Costa A, Navazio L, Szabo I. 2018. The contribution of organelles to plant intracellular calcium signalling. J Exp Bot 10.1093/jxb/ery185.

Ding P, Rekhter D, Ding Y, Feussner K, Busta L, Haroth S, Xu S, Li X, Jetter R, Feussner I, et al. 2016. Characterization of a pipecolic acid biosynthesis pathway required for systemic acquired resistance. Plant Cell 28(10): 2603–2615.

Ding Y, Sun T, Ao K, Peng Y, Zhang Y, Li X, Zhang Y. 2018. Opposite roles of salicylic acid receptors npr1 and npr3/npr4 in transcriptional regulation of plant immunity. Cell 173(6): 1454–1467 e1415.

Dubiella U, Seybold H, Durian G, Komander E, Lassig R, Witte CP, Schulze WX, Romeis T. 2013. Calcium-dependent protein kinase/nadph oxidase activation circuit is required for rapid defense signal propagation. Proc Natl Acad Sci U S A 110(21): 8744–8749.

Durrant WE, Dong X. 2004. Systemic acquired resistance. Annu Rev Phytopathol 42: 185– 209.

Fu ZQ, Dong X. 2013. Systemic acquired resistance: Turning local infection into global defense. Annu Rev Plant Biol 64: 839–863.

Fu ZQ, Yan S, Saleh A, Wang W, Ruble J, Oka N, Mohan R, Spoel SH, Tada Y, Zheng N, et al. 2012. Npr3 and npr4 are receptors for the immune signal salicylic acid in plants. Nature 486(7402): 228–232.

Galon Y, Nave R, Boyce JM, Nachmias D, Knight MR, Fromm H. 2008. Calmodulin-binding transcription activator (camta) 3 mediates biotic defense responses in arabidopsis. FEBS Lett 582(6): 943–948.

Gao X, Chen X, Lin W, Chen S, Lu D, Niu Y, Li L, Cheng C, McCormack M, Sheen J, et al. 2013. Bifurcation of arabidopsis nlr immune signaling via ca^2+^-dependent protein kinases. PLoS Pathog 9(1): e1003127.

Gao X, He P. 2013. Nuclear dynamics of arabidopsis calcium-dependent protein kinases in effector-triggered immunity. Plant Signal Behav 8(4): e23868.

Glazebrook J, Zook M, Mert F, Kagan I, Rogers EE, Crute IR, Holub EB, Hammerschmidt R, Ausubel FM. 1997. Phytoalexin-deficient mutants of arabidopsis reveal that pad4 encodes a regulatory factor and that four pad genes contribute to downy mildew resistance. Genetics 146(1): 381–392.

Glinski M, Romeis T, Witte CP, Wienkoop S, Weckwerth W. 2003. Stable isotope labeling of phosphopeptides for multiparallel kinase target analysis and identification of phosphorylation sites. Rapid Commun Mass Spectrom 17(14): 1579–1584.

Gruner K, Griebel T, Navarova H, Attaran E, Zeier J. 2013. Reprogramming of plants during systemic acquired resistance. Front Plant Sci 4: 252.

Hake K, Romeis T. 2018. Protein kinase-mediated signalling in priming: Immune signal initiation, propagation, and establishment of long-term pathogen resistance in plants. Plant Cell Environ 10.1111/pce.13429.

Hartmann M, Kim D, Bernsdorff F, Ajami-Rashidi Z, Scholten N, Schreiber S, Zeier T, Schuck S, Reichel-Deland V, Zeier J. 2017. Biochemical principles and functional asects of pipecolic acid biosynthesis in plant immunity. Plant Physiol 174(1): 124–153.

Hartmann M, Zeier T, Bernsdorff F, Reichel-Deland V, Kim D, Hohmann M, Scholten N, Schuck S, Brautigam A, Holzel T, et al. 2018. Flavin monooxygenase-generated n-hydroxypipecolic acid is a critical element of plant systemic immunity. Cell 173: 456– 469.

Hilker M, Schwachtje J, Baier M, Balazadeh S, Baurle I, Geiselhardt S, Hincha DK, Kunze R, Mueller-Roeber B, Rillig MC, et al. 2015. Priming and memory of stress responses in organisms lacking a nervous system. Biol Rev Camb Philos Soc 20(10): 12215.

Kadota Y, Sklenar J, Derbyshire P, Stransfeld L, Asai S, Ntoukakis V, Jones JD, Shirasu K, Menke F, Jones A, et al. 2014. Direct regulation of the nadph oxidase rbohd by the prr-associated kinase bik1 during plant immunity. Mol Cell 54(1): 43–55.

Kobayashi M, Ohura I, Kawakita K, Yokota N, Fujiwara M, Shimamoto K, Doke N, Yoshioka H. 2007. Calcium-dependent protein kinases regulate the production of reactive oxygen species by potato nadph oxidase. Plant Cell 19(3): 1065–1080.

Kobayashi M, Yoshioka M, Asai S, Nomura H, Kuchimura K, Mori H, Doke N, Yoshioka H. 2012. Stcdpk5 confers resistance to late blight pathogen but increases susceptibility to early blight pathogen in potato via reactive oxygen species burst. New Phytol 196(1): 223–237.

Liese A, Romeis T. 2013. Biochemical regulation of in vivo function of plant calcium-dependent protein kinases (cdpk). Biochim Biophys Acta 1833(7): 1582–1589.

Liu N, Hake K, Wang W, Zhao T, Romeis T, Tang D. 2017. Calcium-dependent protein kinase5 associates with the truncated nlr protein tir-nbs2 to contribute to exo70b1-mediated immunity. Plant Cell 29(4): 746–759.

Manohar M, Tian M, Moreau M, Park SW, Choi HW, Fei Z, Friso G, Asif M, Manosalva P, von Dahl CC, et al. 2015. Identification of multiple salicylic acid-binding proteins using two high throughput screens. Front Plant Sci 5(777).

Martinez-Medina A, Flors V, Heil M, Mauch-Mani B, Pieterse CMJ, Pozo MJ, Ton J, van Dam NM, Conrath U. 2016. Recognizing plant defense priming. Trends Plant Sci 21(10): 818–822.

Mishina TE, Zeier J. 2006. The arabidopsis flavin-dependent monooxygenase fmo1 is an essential component of biologically induced systemic acquired resistance. Plant Physiol 141(4): 1666–1675.

Monaghan J, Matschi S, Romeis T, Zipfel C. 2015. The calcium-dependent protein kinase cpk28 negatively regulates the bik1-mediated pamp-induced calcium burst. Plant Signal Behav 10(5): e1018497.

Monaghan J, Matschi S, Shorinola O, Rovenich H, Matei A, Segonzac C, Malinovsky FG, Rathjen JP, MacLean D, Romeis T, et al. 2014. The calcium-dependent protein kinase cpk28 buffers plant immunity and regulates bik1 turnover. Cell Host Microbe 16(5): 605–615.

Mori IC, Murata Y, Yang Y, Munemasa S, Wang YF, Andreoli S, Tiriac H, Alonso JM, Harper JF, Ecker JR, et al. 2006. Cdpks cpk6 and cpk3 function in aba regulation of guard cell s-type anion- and ca^2+^-permeable channels and stomatal closure. PLoS Biol 4(10): e327.

Návarová H, Bernsdorff F, Doring AC, Zeier J. 2012. Pipecolic acid, an endogenous mediator of defense amplification and priming, is a critical regulator of inducible plant immunity. Plant Cell 24(12): 5123–5141.

Reimer-Michalski EM, Conrath U. 2016. Innate immune memory in plants. Semin Immunol 28(4): 319–327.

Rietz S, Stamm A, Malonek S, Wagner S, Becker D, Medina-Escobar N, Vlot AC, Feys BJ, Niefind K, Parker JE. 2011. Different roles of enhanced disease susceptibility1 (eds1) bound to and dissociated from phytoalexin deficient4 (pad4) in arabidopsis immunity. New Phytol 191(1): 107–119.

Romeis T, Piedras P, Jones JD. 2000. Resistance gene-dependent activation of a calcium-dependent protein kinase in the plant defense response. Plant Cell 12(5): 803–816.

Schulz P, Herde M, Romeis T. 2013. Calcium-dependent protein kinases: Hubs in plant stress signaling and development. Plant Physiol 163(2): 523–530.

Serrano M, Wang B, Aryal B, Garcion C, Abou-Mansour E, Heck S, Geisler M, Mauch F, Nawrath C, Metraux JP. 2013. Export of salicylic acid from the chloroplast requires the multidrug and toxin extrusion-like transporter eds5. Plant Physiol 162(4): 1815–1821.

Seybold H, Trempel F, Ranf S, Scheel D, Romeis T, Lee J. 2014. Ca^2+^ signalling in plant immune response: From pattern recognition receptors to ca^2+^ decoding mechanisms. New Phytol 204(4): 782–790.

Shah J, Zeier J. 2013. Long-distance communication and signal amplification in systemic acquired resistance. Front Plant Sci 4: 30.

Stael S, Wurzinger B, Mair A, Mehlmer N, Vothknecht UC, Teige M. 2012. Plant organellar calcium signalling: An emerging field. J Exp Bot 63(4): 1525–1542.

Sun T, Busta L, Zhang Q, Ding P, Jetter R, Zhang Y. 2018. Tgacg-binding factor 1 (tga1) and tga4 regulate salicylic acid and pipecolic acid biosynthesis by modulating the expression of systemic acquired resistance deficient 1 (sard1) and calmodulin-binding protein 60g (cbp60g). New Phytol 217(1): 344–354.

Sun T, Zhang Y, Li Y, Zhang Q, Ding Y, Zhang Y. 2015. Chip-seq reveals broad roles of sard1 and cbp60g in regulating plant immunity. Nat Commun 6: 10159.

Truman W, Glazebrook J. 2012. Co-expression analysis identifies putative targets for cbp60g and sard1 regulation. BMC Plant Biol 12: 216.

Wang J, Grubb LE, Wang J, Liang X, Li L, Gao C, Ma M, Feng F, Li M, Li L, et al. 2018. A regulatory module controlling homeostasis of a plant immune kinase. Mol Cell 69(3): 493–504 e496.

Wang L, Tsuda K, Sato M, Cohen JD, Katagiri F, Glazebrook J. 2009. Arabidopsis cam binding protein cbp60g contributes to mamp-induced sa accumulation and is involved in disease resistance against pseudomonas syringae. PLoS Pathog 5(2): e1000301.

Wang L, Tsuda K, Truman W, Sato M, Nguyen le V, Katagiri F, Glazebrook J. 2011. Cbp60g and sard1 play partially redundant critical roles in salicylic acid signaling. Plant J 67(6): 1029–1041.

Wang Y, Schuck S, Wu J, Yang P, Doring AC, Zeier J, Tsuda K. 2018. A mpk3/6-wrky33-ald1-pipecolic acid regulatory loop contributes to systemic acquired resistance. Plant Cell 30(10): 2480–2494.

Wiermer M, Feys BJ, Parker JE. 2005. Plant immunity: The eds1 regulatory node. Curr Opin Plant Biol 8(4): 383–389.

Wildermuth MC, Dewdney J, Wu G, Ausubel FM. 2001. Isochorismate synthase is required to synthesize salicylic acid for plant defence. Nature 414(6863): 562–565.

Wu Y, Zhang D, Chu JY, Boyle P, Wang Y, Brindle ID, De Luca V, Despres C. 2012. The arabidopsis npr1 protein is a receptor for the plant defense hormone salicylic acid. Cell Rep 1(6): 639–647.

Yan S, Dong X. 2014. Perception of the plant immune signal salicylic acid. Curr Opin Plant Biol 20: 64–68.

Ye W, Muroyama D, Munemasa S, Nakamura Y, Mori IC, Murata Y. 2013. Calcium-dependent protein kinase cpk6 positively functions in induction by yeast elicitor of stomatal closure and inhibition by yeast elicitor of light-induced stomatal opening in arabidopsis. Plant Physiol 163(2): 591–599.

Zhang Y, Xu S, Ding P, Wang D, Cheng YT, He J, Gao M, Xu F, Li Y, Zhu Z, et al. 2010. Control of salicylic acid synthesis and systemic acquired resistance by two members of a plant-specific family of transcription factors. Proc Natl Acad Sci U S A 107(42): 18220–18225.

Zheng XY, Zhou M, Yoo H, Pruneda-Paz JL, Spivey NW, Kay SA, Dong X. 2015. Spatial and temporal regulation of biosynthesis of the plant immune signal salicylic acid. Proc Natl Acad Sci U S A 112(30): 9166–9173.

Zhou M, Lu Y, Bethke G, Harrison BT, Hatsugai N, Katagiri F, Glazebrook J. 2018. Wrky70 prevents axenic activation of plant immunity by direct repression of sard1. New Phytol 217(2): 700–712.

